# *Wolbachia*-mediated antiviral protection is driven by its multimodal effects on *Drosophila melanogaster* metabolism

**DOI:** 10.1101/2025.10.22.681277

**Authors:** Sarah M. Walsh, Tyson G. Thomson, Karyn N. Johnson, Jeremy C. Brownlie, Horst Joachim Schirra

## Abstract

*Wolbachia*, a common endosymbiont of *Drosophila melanogaster*, provides a profound antiviral effect in insects. *Wolbachia*-mediated viral interference has been employed to limit the spread of arboviruses, including dengue. However, the mechanisms underpinning *Wolbachia*-mediated viral interference are not consistently understood. Previous studies have identified resource competition as a potential mechanism by which *Wolbachia* disrupts viral replication. Our study uses nuclear magnetic resonance (NMR)-based metabolomics to characterize the bi- and tripartite host-*Wolbachia*-virus interactions using the model insect *Drosophila melanogaster*, the protective *Wolbachia* strain *w*Mel and the pathogenic Drosophila C virus (DCV). The findings reveal that *w*Mel-infected flies showed increased simple carbohydrate catabolism relative to uninfected *Drosophila.* DCV infection perturbed nucleotide synthesis and nucleotide abundance in *Drosophila* compared to uninfected *Drosophila*, driving metabolism to likely meet the viral replication demands imposed on the host. Notably, co-infected *Drosophila* exhibited a metabolic profile more similar to *w*Mel-infected flies than DCV-infected flies, suggesting that *w*Mel drives metabolism in a direction that at least temporarily inhibits DCV replication. The metabolic profile is indicative of a hypoxic environment that has been known to trigger immune pathways that further contribute to *Wolbachia*-mediated pathogen blocking. It is probable that *Wolbachia*-mediated antiviral protection is a multimodal consequence of *w*Mel’s influence on host metabolism, rather than a single mechanism. These findings will guide future research and contribute to the continued success of *Wolbachia*-based vector control strategies against RNA arboviruses, potentially leading to novel approaches for defending against such pathogens and improving vector control strategies.

**Significance Statement:** *Wolbachia*-mediated antiviral protection has been used extensively to control the spread of insect-borne viruses, with the release of *Wolbachia*-infected mosquitoes eradicating local dengue transmission in some regions. Despite this real-world impact, how *Wolbachia* disrupted viruses remained unclear, with three common mechanisms predicted: (1) competition between *Wolbachia* and virus for host metabolites, or stimulation of host antiviral pathways via (2) immune priming and/or (3) the RNA interference pathway. Our findings reveal that in *Drosophila melanogaster*, *Wolbachia* does compete for some metabolites, but it exerts a greater effect by altering host metabolism which in turn creates an intracellular environment unfavorable for viral replication and indirectly triggers an immune response. Thus, the mechanism by which *Wolbachia* disrupts viruses relies on both metabolic and immune pathways. Understanding these metabolic mechanisms is crucial for optimizing and sustaining this vector control strategy and for shaping future efforts to combat vector-borne diseases.

## Introduction

*Wolbachia* are maternally transmitted gram-negative obligate endosymbionts that infect an estimated 50% of terrestrial arthropods including, insects, arachnids, isopods and filarial nematodes [1]. Some strains of *Wolbachia* that infect insects form more complex and varied partnerships with their hosts. All strains are considered reproductive parasites, as nearly all manipulate host reproductive or sex determination systems to increase their frequency within a host [2, 3]. For some strains, *Wolbachia* provides a fitness benefit to their host including metabolic provisioning [4, 5] or protection against viral pathogens [6, 7]. One such example is *w*Mel, which naturally infects *Drosophila melanogaster,* imposes cytoplasmic incompatibility [2, 8], modulates iron-metabolism to increase female fecundity [9] and can reduce the pathogenicity of RNA viruses and delay the accumulation of virus particles in the host [6, 7].

*Wolbachia*-mediated antiviral protection was initially discovered in *D. melanogaster* [6, 7] but has since been observed for *Wolbachia* strains that infect other insect species [6, 10–16]. *w*Mel mediated virus interference is maintained in novel host backgrounds [11], including the major mosquito vector of dengue fever *Aedes aegypti* [12, 13, 17, 18] and planthoppers [19]. As a result of *w*Mel-infected mosquitoes being released into wild populations around the world, a significant reduction in prevalence and in some instances complete eradication of dengue from communities has been achieved [20–22]. While *Wolbachia*-mediated viral interference has been described in natural systems for more than a decade and used as a biocontrol agent, the detailed mechanism(s) of interference remain to be defined.

There are several mechanisms that may contribute to *Wolbachia*-mediated antiviral protection, but these mechanisms frequently fail to completely explain this phenomenon. These mechanisms include RNA interference, antiviral immune responses, cellular tropism, and metabolic interaction. While *Wolbachia* interacts with the RNA interference pathway in insects [23], studies suggest that *Wolbachia*-mediated antiviral protection works independently of these systems [24]. Similarly, while some *Wolbachia* infections stimulate the host immune system, these are typically limited to novel host infections or do not correlate with *Wolbachia* mediated antiviral protection [25, 26]. *Wolbachia*-mediated antiviral protection is an intracellular response [27]. *Wolbachia*-free cells when exposed to an RNA virus will fail to inhibit viral infection within the host; however, *Wolbachia*-infected cells will inhibit replication of the virus in that cell [27, 28]. This mechanism therefore reveals that *Wolbachia*’s pathogen blocking is not a systemic process, but rather localized to the endosymbiont. *Wolbachia* and viruses are both metabolically dependent upon their insect hosts to complete their lifecycles, providing an opportunity for symbiont-conflicts to arise. For example, cholesterol is a limiting metabolite for virus replication and *Wolbachia* proliferation [29]. An artificial increase of cholesterol in insect diets correlated with an increase in DCV accumulation and decreased pathogen blocking in *w*Mel-infected *Drosophila* [30, 31]. Furthermore, *Wolbachia* perturbs other lipid classes, such as an increase in fatty acids as acylcarnitines are catabolized, causing a depletion in ATP production crucial for viral replication [32]. A second class of metabolites that are correlated with *Wolbachia*-mediated antiviral protection are nucleotides. In *D. melanogaster*, *w*Mel infection is correlated with increased *prat2* gene expression, involved in the *de novo* synthesis of purines, which correlated with *Wolbachia*’s ability to suppress Sindbus virus [33]. Lindsey et al. (2021) proposed that there were, yet unclear, downstream consequences for example increased activity of the purine salvage pathway that negatively effects the cellular processes of Sindbis virus [33].

Taken together, this evidence points towards metabolic interaction as a fundamental mechanism for *Wolbachia*-mediated pathogen blocking. While metabolic competition for lipids appears to underpin *Wolbachia*-mediated viral interference it is unclear whether other metabolites, such as amino acids, could also contribute to this phenotype. Here, we have applied nuclear magnetic resonance (NMR)-based metabolomics to take a non-hypothesis driven approach to identify points of metabolic interaction among *Drosophila melanogaster, w*Mel and DCV.

## Results

### Confirmation of the effect of a Drosophila C virus in *D. melanogaster*

To confirm the approximate time points of 50 % survival for *w*Mel and *w*Mel-free Fly lines infected with Drosophila C virus (DCV) a survival assay was performed. In *w*Mel-free *Drosophila,* DCV induced 100% mortality within 9 days post infection (dpi), while *w*Mel-infected *Drosophila* survived until 19 dpi (Supplementary Fig. S1A). Approximately 50 % survival for *w*Mel-free and *w*Mel-infected *Drosophila* lines were observed at 5 and 9 dpi, respectively (Supplementary Fig. S1A). These time points were used for all subsequent experiments (Fig. 1A). It was re-confirmed, in agreement with previous research [6, 7], that this delay in DCV-induced mortality corresponded to a delay in viral accumulation (i.e., *w*Mel delays both DCV accumulation, and virus-induced death) (Supplementary Fig. S1B).

**Figure 1.**
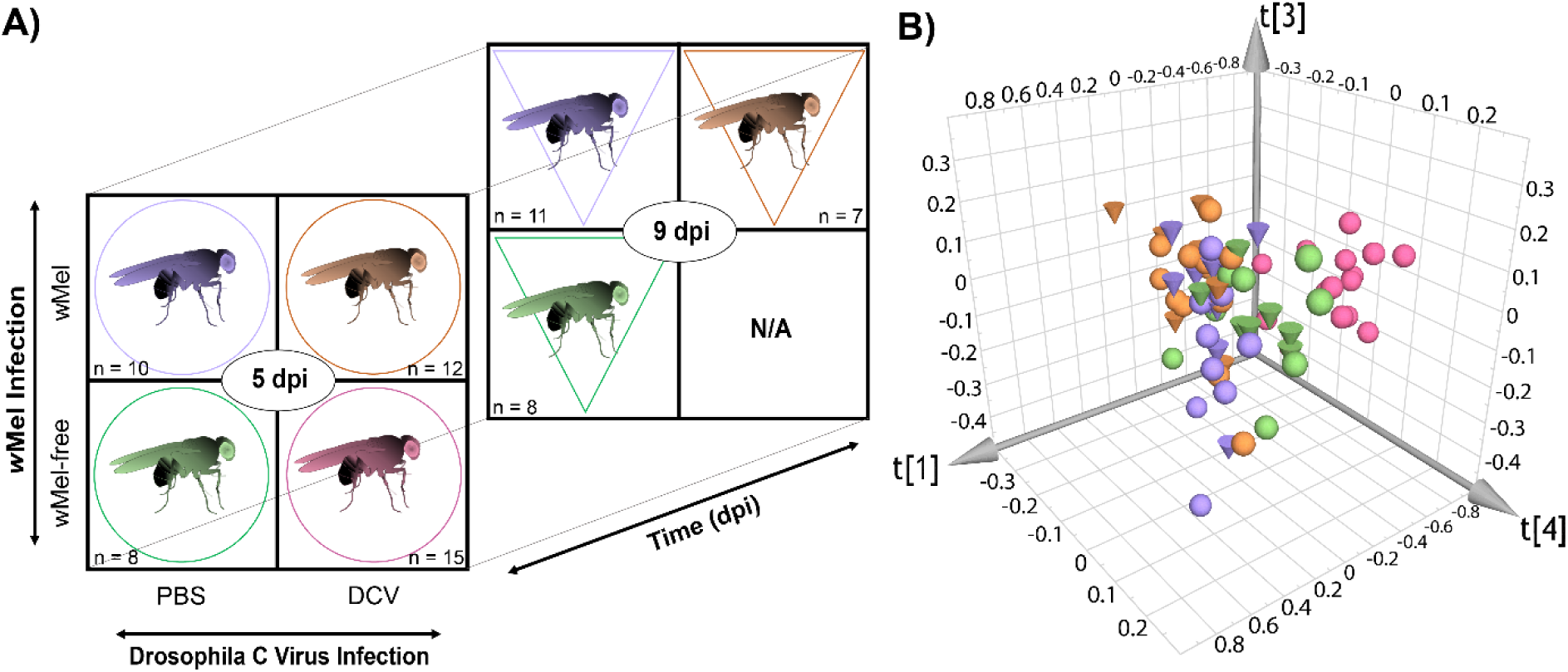
Global multivariate statistical analysis reveals the study cohorts are metabolically unique A) Seven Drosophila cohorts—uninfected (green), DCV-infected (pink), *w*Mel-infected (purple), and co-infected (orange)—were analyzed for metabolic differences at 5 dpi (circles) and 9 dpi (inverted triangles). Cohort sizes are denoted by *n*. B) PCA scores plot of all sample cohorts (Supplementary Table S1: model **M2**). The principal component 1 on three-dimensional (3D) is indicated by t[1], principal component 4 by t[4], and finally principal component 3 is t[3].

### Global Multivariate Analysis: *w*Mel-, DCV- and co-infected *Drosophila* have unique metabolic profiles

Eighty-four *Drosophila* samples were extracted, and the water-soluble extracts collected. Following initial NMR analysis, 11 samples were excluded due to poor water suppression in the NMR spectra and not analyzed further. Multivariate statistical analysis (MVSA) was applied to 73 samples in seven cohorts (Fig 1B): co-infected at 5 dpi (*n* = 12); DCV-infected at 5 dpi (*n* = 15); *w*Mel-infected at 5 dpi (*n* = 10); uninfected at 5 dpi (*n* = 8); co-infected at 9 dpi (*n* = 7); *w*Mel-infected at 9 dpi (*n* = 11), and uninfected at 9 dpi (*n* = 8). As DCV had killed more than 50% of *w*Mel-free DCV-infected *Drosophila* at 9 dpi, samples could not be collected for this cohort (Supplementary Fig. S1).

A multivariate statistical analysis of all samples (model **M2**; Fig. 1B) identified systematic global differences among the metabolic profiles of the cohorts (Figures of merit for all models in Supplementary Table S1). The DCV-infected cohort had the greatest global metabolic difference compared to all other cohorts. Comparing cohorts across time (i.e., 5 and 9 dpi) showed that uninfected, single *w*Mel-infected, and co-infected cohorts clustered more closely together, but were still distinctly separate. Thus, DCV, *w*Mel, or co-infection of both microbes uniquely affect *Drosophila* global metabolism (Fig. 1B). To study those effects in more detail, a series of PLS models of pairwise comparisons of two study groups were created and interpreted.

### The effect of time on the metabolism of *Drosophila*

To examine the effect of time on the metabolism of *Drosophila*, uninfected cohorts at 5 and 9 dpi were compared with PLS (model **M16**). The model showed that there was a difference between the metabolic profiles of uninfected *D. melanogaster* at 5 and 9 dpi, revealing that the two cohorts mostly clustered separately (Supplementary Fig. S2A).

In the corresponding bivariate loadings plot (Supplementary Fig. 2B) these differences were highlighted. To confirm the validity of these metabolic differences, a univariate analysis was conducted (Supplementary Table S2 and Fig. S3). Both analyses showed that many metabolites were at higher concentrations at 5 dpi relative to 9 dpi. These included inosine, glucose, maltose, lysine, propionate, and arginine. Overall, the metabolic profile of uninfected *Drosophila* reflected a hypermetabolic state, with higher levels of amino acids and carbon sources relative to uninfected *Drosophila* at 9 dpi.

### The effect of *w*Mel infection on the metabolism of *Drosophila*

Model **M2** (Fig. 1B) indicated that *w*Mel infection clustered uniquely in the global PCA analysis. PLS models showed that there was a difference between the metabolic profiles of *w*Mel-infected *D. melanogaster* and *w*Mel*-*free *D. melanogaster* at 5 dpi (Fig. 2D, model **M17**) and at 9 dpi (Supplementary Fig. S4, model **M18**), consistent with a previous study [34]. These metabolic differences were confirmed by the scores plots, (Fig. 2A and Supplementary Fig. S4B, respectively), while both multivariate and univariate analyses of the *w*Mel-infected cohort at 5 dpi with the *w*Mel-free cohort at 5 dpi (Fig. 2D, Supplementary Fig. S3, and Supplementary Table S2) showed an overall increase in glycolytic end products, amino acids and carbohydrates in *w*Mel-infected *Drosophila* relative to *w*Mel-free *Drosophila.* The analyses showed that *w*Mel-infected *Drosophila* had higher levels of adenosine, gluconate, proline, and tyrosine, compared to *w*Mel-free *Drosophila*. *w*Mel-infected *Drosophila* had lower levels of pyruvate, glucose, maltose, lysine, alanine, arginine, and 2-oxogluatarate, compared to *w*Mel-free *Drosophila*. When comparing the *w*Mel-infected cohort at 9 dpi with the *w*Mel-free cohort at 9 dpi the multivariate and univariate analyses showed that *w*Mel-infected *Drosophila* at 9 dpi (Supplementary Fig. S7, Fig. S5, and Table S9) had different metabolic levels compared to 5 dpi, most of the metabolites that were at higher concentrations in the uninfected cohort at 5 dpi relative to the *w*Mel-infected cohort were found to be at lower levels at 9 dpi. However, adenosine, gluconate, proline, and tyrosine were found to be similarly higher levels in *w*Mel infection relative to the uninfected cohort at 9 dpi. Therefore, it is likely that time post injection affects the metabolic profile, and therefore the reason there are changes between 5 and 9 dpi comparing *Wolbachia* infection to uninfected insects. At 9 dpi the *w*Mel*-*infected cohort relative to uninfected *Drosophila* (Supplementary Fig. S4), had a significant increase in glycine, lactate, acetate, valine, proline and O-phosphocholine. Overall, there was a primary increase in glycolytic end products, amino acids and carbohydrates.

**Figure 2.**
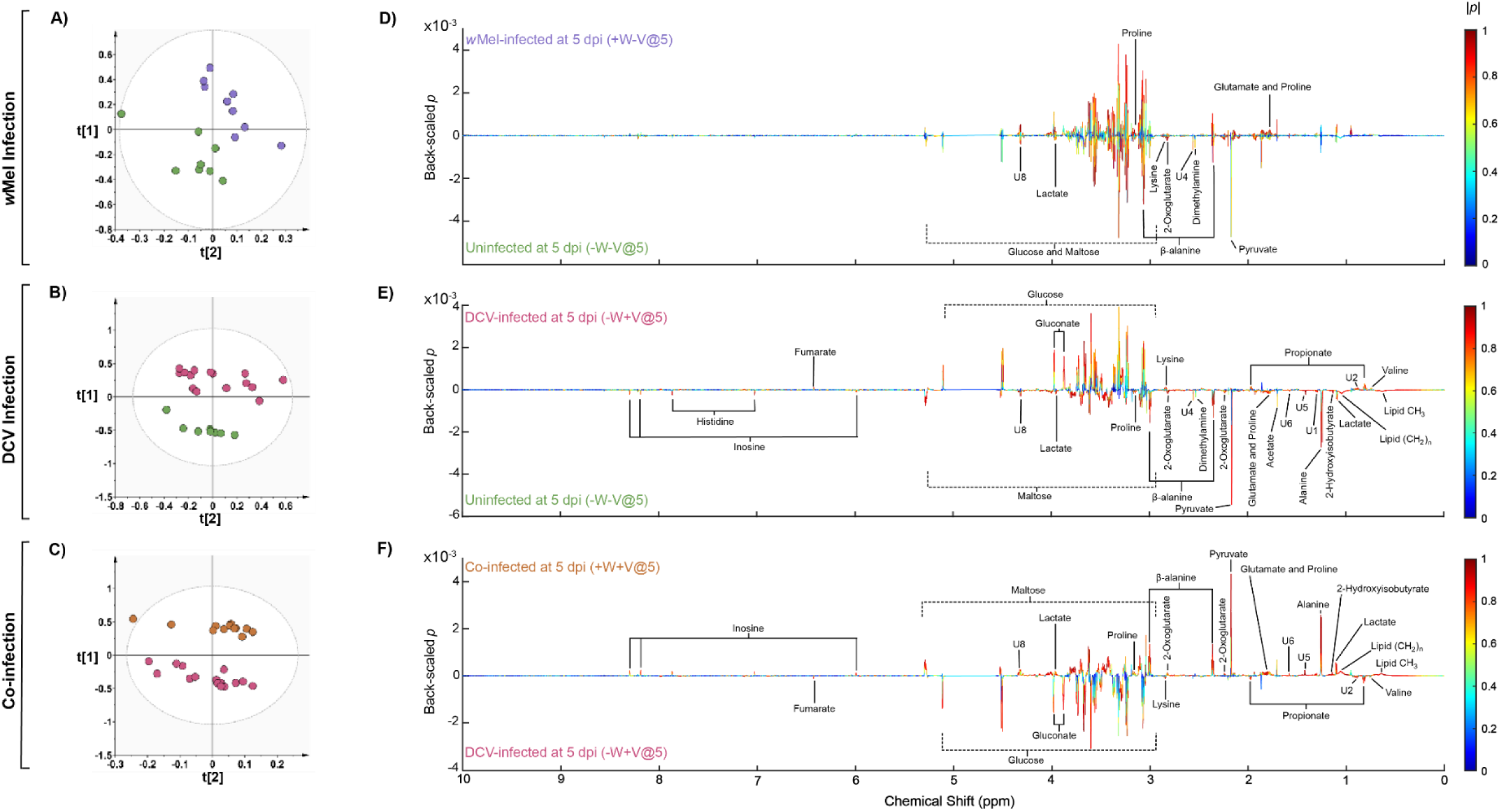
Pairwise comparisons examining *w*Mel infection, DCV infection, and co-infected *Drosophila* against DCV infection. A-C) PLS scores plots comparing metabolic profiles of *Drosophila* cohorts at 5 dpi: *w*Mel-infected versus uninfected (+W−V@5 [purple] vs. −W−V@5 [green]), DCV-infected versus uninfected (−W+V@5 [pink] vs. −W−V@5 [green]), and co-infected versus DCV-infected (+W+V@5 [orange] vs. −W+V@5 [pink]). The x and y axes represent components t[1] and t[2] respectively. D-F) One-dimensional (1D) bivariate loadings plots of the corresponding pairwise PLS models. The back-scaled loadings coefficients *p* were plotted against the chemical shift (ppm) for the bucketed multivariate statistical analysis *X*-matrices’ variables obtained from the 1D 1H NMR spectra. Also, the values of the correlation coefficients |*p*(*corr*)| were superimposed on the bivariate loadings plot as a heatmap color scale. Metabolites that were significantly altered between comparison were annotated in black (|*p*|≥ 0.015; |*p*(*corr*)| ≥ 0.6), and VIP ≥ 1).

### The effect of DCV infection on the metabolism of *Drosophila*

Both global PCA and PLS pairwise model (M19) showed that DCV infection was a latent variable separating the singly DCV-infected cohort from the other six cohorts (Fig. 2B).

The bivariate loadings plot (Fig. 2E) revealed that DCV-infected *Drosophila* had higher levels of fumarate, glucose, gluconate, lysine, propionate, and valine relative to uninfected *Drosophila*. The loadings plot also revealed lower levels of histidine, inosine, maltose, lactate, proline, β-alanine, 2-oxoglutarate, dimethylamine, pyruvate, glutamate, acetate, alanine, 2-hydroxyisobutyrate, and lipids. These relative changing metabolite levels in metabolite concentration were confirmed through further univariate analysis (Supplementary Fig. S3 and Table S2).

### The effect of *w*Mel infection on the metabolism of DCV-infected *Drosophila*

Metabolic profiles of *Drosophila* infected by both *w*Mel and DCV, either 5 or 9 dpi, were different to 5 dpi *w*Mel-free DCV-infected *Drosophila* cohort (PLS model **M20**; scores plot Fig. 2C). The corresponding loadings plot (Fig. 2F) showed points of competition between *w*Mel and DCV and metabolic trends indicative of a hypoxic environment. The loadings plot (Fig. 2F) revealed that co-infected *Drosophila* had lower levels of fumarate, gluconate, glucose, propionate, valine, and lysine relative to *w*Mel-free DCV-infected *Drosophila*. There were also higher levels of inosine, maltose, lactate, proline, beta-alanine, pyruvate, alanine, glutamate, and lipids relative to 5 dpi *w*Mel-free DCV-infected *Drosophila* (Fig. 2F). Relative changes in metabolite concentration were confirmed by univariate analysis (Supplementary Fig. S3 and Table S2).

Pairwise comparisons between co-infected and *w*Mel-infected DCV-free *D. melanogaster* at 5 and 9 dpi (**M10** and **M11**) revealed that there were minimal metabolic differences between co-infected cohorts to DCV-free *w*Mel-infected cohorts. This was further supported by the univariate analysis that found that there were no significant changes in metabolite concentrations for single *w*Mel infection relative to co-infected *Drosophila* at 5 or 9 dpi (Supplementary Fig. S3 and Table S2). However, at 9 dpi, in the PCA of all *w*Mel-infected cohorts revealed that the co-infected cohort began to separate away from the other *w*Mel-infected cohorts in the scores plot (Fig. 3C). Metabolically the profile of the *w*Mel-infected *Drosophila* is similar, even co-infected with DCV, until it we assume that DCV infection likely begins to overwhelm the system.

**Figure 3.**
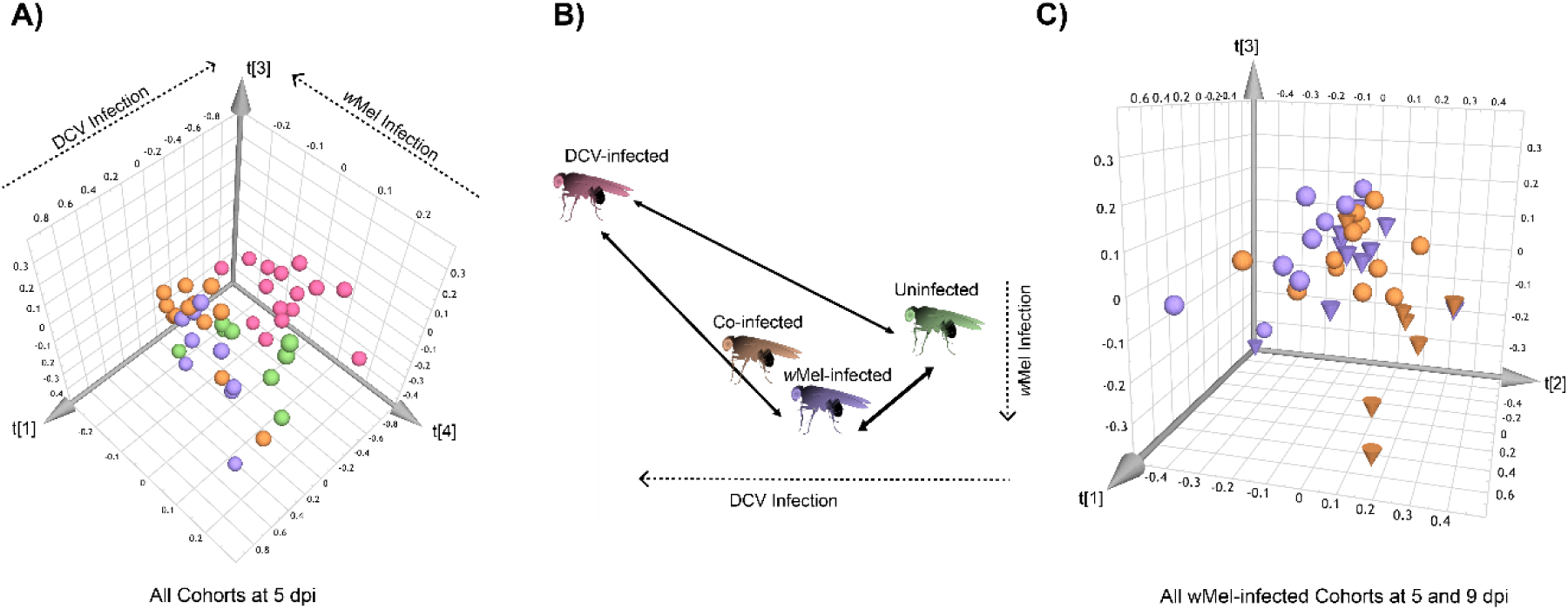
*w*Mel temporarily delays the metabolic demands of DCV on *Drosophila*. A) Four-way PCA score plot of all cohorts at 5 dpi (model **M14**), with principal components t[1], t[4], and t[3] on the x-, y-, and z-axes, respectively. B) Depiction of metabolic similarity between cohorts at 5 dpi. The line distances in the diagram (B) represent are arbitrary score of similarity seen in (A), where the short the line the more similar the cohorts are metabolically. C) PCA analysis of all *w*Mel-infected cohorts (**M15**), with principal components t[1], t[2], and t[3] on the x-, y-, and z-axes, respectively. Colors and shapes match previous figures: spheres represent 5 dpi; cones 9 dpi; green indicates uninfected, pink DCV-infected, purple *w*Mel-infected, and orange co-infected cohorts.

To determine points of metabolic competition between *Wolbachia* and DCV, a four-way comparison using PCA (model **M14**) was conducted. All four cohorts (co-infected *Drosophila*, *w*Mel-free DCV-infected *Drosophila*, *w*Mel-infected *Drosophila*, and uninfected *Drosophila* at 5 dpi) clustered into unique groups, emphasizing the differences in metabolic profiles between them (Fig. 3A). However, 5 dpi co-infected *Drosophila* are more metabolically similar to both 5 and 9 dpi *w*Mel-infected *Drosophila* as compared to 5 dpi DCV-infected *Drosophila* even though both cohorts are infected with DCV and are the same time points post infection (Fig. 3B). The shared metabolic profile of co-infected *Drosophila* at 5 dpi with *w*Mel-infected *Drosophila* cohorts (Fig. 3C) correlates with the protective effect *Wolbachia* provides against DCV accumulation when injected with the virus and delayed pathology (Supplementary Fig. S1). However, by 9 dpi, the metabolic profile of the co-infected cohort has diverged from the 5 and 9 dpi *w*Mel infected cohorts but does not yet metabolically resemble the DCV-infected *Drosophila* (Fig. 3C).

## Discussion

Both microbes in this study infect the host intracellularly and metabolically depend on the host to survive. *w*Mel is a maternally inherited endosymbiont whose fitness (i.e., the ability to be passed onto the next generation) depends on its host. For *w*Mel to be transmitted to the next generation the host must survive long enough to produce offspring. DCV is a vertically and horizontally transmitted pathogen that needs its host to survive long enough for it to replicate. It is this fundamental difference in biology that we believe creates diverging and competing metabolic host-guest interactions between these different microbes. Together, both DCV and *w*Mel have the potential to drive the host’s metabolic state to reflect their respective metabolic demands as they interact within the host’s cells as the microbes replicate. It has been widely hypothesized that metabolic competition is likely behind *Wolbachia*-mediated antiviral protection [31, 35, 36]. In this study the metabolic changes caused by *w*Mel, Drosophila C virus (DCV), and co-infections in *Drosophila melanogaster* were characterized, to understand the metabolic tripartite interaction behind *Wolbachia*-mediated antiviral protection.

We observed metabolic changes in *Wolbachia* infection characteristic of hypoxia including increased glycolytic activity and decreased TCA cycle activity [34, 37] suggesting that *w*Mel competes with the host for oxygen [34, 38]. Hypoxia is known to lead to the accumulation of reactive oxygen species (ROS) and consequentially oxidative stress. Importantly, *w*Mel infection is known to correlate with an increase in ROS and oxidative stress [39]. ROS can act as a signaling molecule, in which case, an upregulation of ROS promotes the initiation of signaling pathways. For example, one of the pathways indirectly initiated by ROS in *Drosophila* is the extracellular signal-regulated kinase (ERK) signaling cascade [40]. The ERK pathway has proven critical in antiviral resistance in *Drosophila*, where mutant loss-of-function ERK *Drosophila* were more susceptible to virus-induced mortality [41]. This is important as *w*Mel in DCV-infected *Drosophila* also generates a hypoxic environment (Fig. 4). However, this ERK studies revealed that while ERK activation is increased in *Wolbachia*-infected species, *Wolbachia*-mediated antiviral protection occurs in absence of this pathway [41]. Indeed, evidence has shown that *w*Mel does not directly stimulate the immune system. Thus, it is more likely that this production of ROS is a consequence of the demands of *w*Mel-infection contributing to *Wolbachia*-mediated antiviral protection by prompting an immune response. Additionally, nonprotective *Wolbachia* strains, such as *w*No and *w*Ha of *Drosophila simulans*, were found to have no change in ROS and oxidative stress [39]. It is possible that non-protective strains do not have the same metabolic demands as that of the protective strains, like *w*Mel, and thus the reason why these nonprotective strains do not induce ROS and therefore cannot protect against viruses.

**Figure 4.**
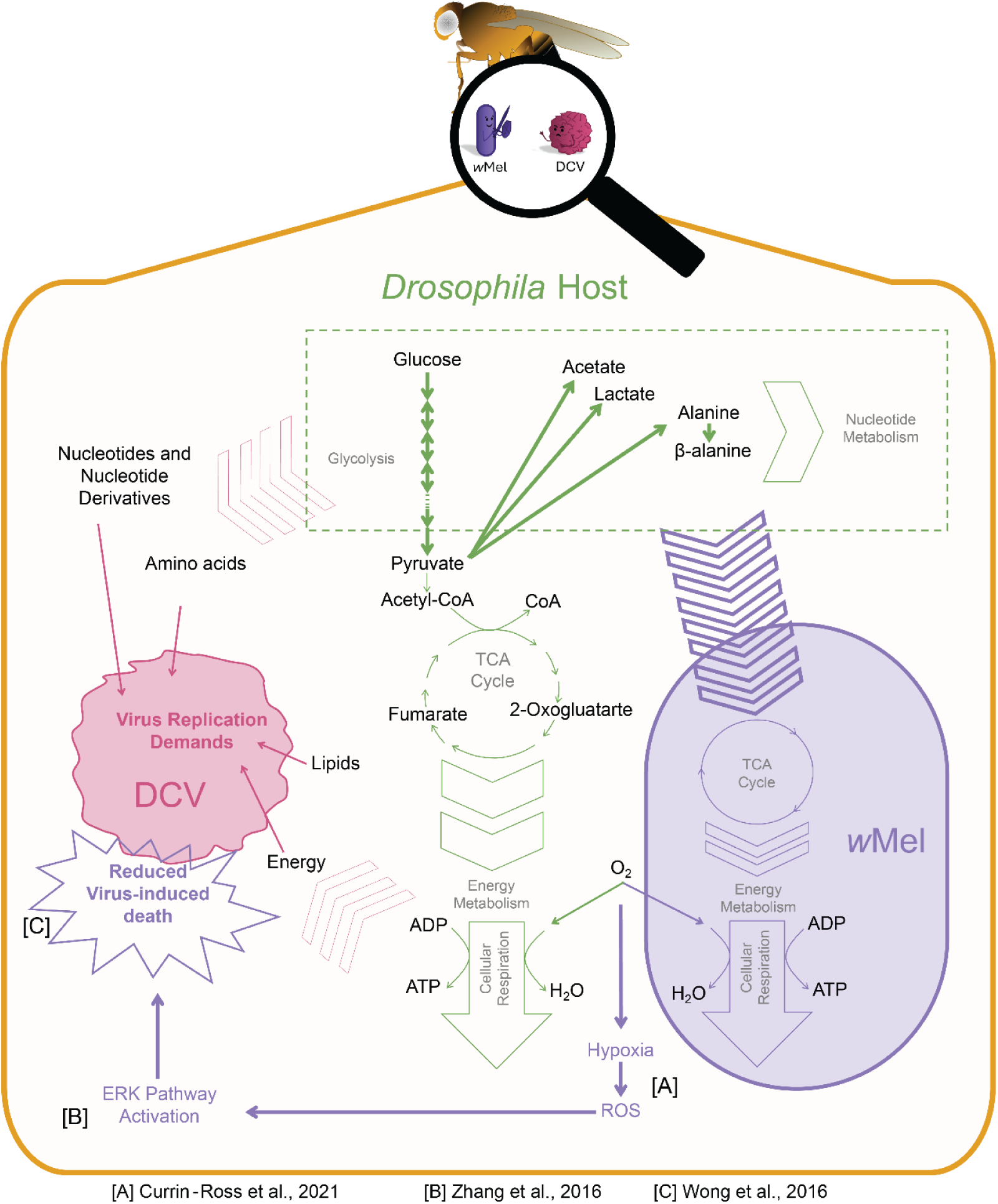
The Metabolic effect of *w*Mel infection in DCV-infected *D. melanogaster*. *w*Mel infection in DCV-infected *D. melanogaster* is associated with an increase in glycolysis and decrease in TCA cycle activity. This in turn leads to the competition of oxygen within the system, inducing hypoxia and ROS within host cells. ROS activates immune signaling pathways such as EKR, which reduces host susceptibility to DCV. However, while TCA cycle activity is decreased, glycolysis is increased, where glycolytic products are increased as a result. Moreover, as a consequence of *w*Mel infection, the metabolic demands for DCV replication demands are not met as metabolites are diverted to *w*Mel. Where the green arrows indicate normal host metabolic behavior; pink arrows indicate DCV’s metabolic demands imposed on the host, and purple arrows indicate *w*Mel’s metabolic processes, demands, and effect. The thicker arrows indicate greater activity of metabolic pathways/effect driven by *w*Mel in the DCV-infected host relative to the *w*Mel-free DCV-infected host.

The metabolic differences between *w*Mel-infected *Drosophila* and *w*Mel-free *Drosophila* reveal further insights into host-symbiont-pathogen interactions. At 5 dpi, *w*Mel-infected *Drosophila* exhibited decreased levels of glycolytic intermediates and end products, carbohydrates, amino acids, and tricarboxylic end products compared to their uninfected counterparts (Fig. 2D).

Notably, in this comparison metabolites such as adenosine, proline, tyrosine, and uracil, were elevated in *w*Mel-infected *Drosophila*. However, by 9 dpi, many of the metabolites initially at lower levels were found at higher levels in the *w*Mel-infected *Drosophila* (Supplementary Fig.S4, Fig. S3, and Table S2). These differences between comparisons at 5 and 9 dpi may reflect metabolic processes associated with wound repair following injection. A previous metabolomic study of *w*Mel in *Drosophila* observed decrease in proline when compared to *w*Mel-free *Drosophila*; it was postulated that increased proline catabolism would increase ROS production [34]. This observation differs from our study which revealed that proline levels were elevated in *w*Mel-infected *Drosophila* [34]. These conflicting results could be attributed to a different genetic background of the *Drosophila*.

In direct contrast to *Wolbachia*, DCV is pathogenic parasitic symbiont, hijacking the host’s cellular machinery to replicate at the expense of host fitness (Fig. 4). Here we observed an increase of monosaccharides and a decrease in disaccharide levels and amino acids relative to uninfected *Drosophila* (Fig. 2E), with a general increase of energy metabolism, through glycolysis and the TCA cycle pathways. While this study is the first metabolomic analysis of DCV, upregulation of glycolysis and glucose metabolism is a common consequence of arboviral infection. Cricket paralysis virus infection of *Drosophila* S2 cells correlated with an increase in carbohydrates [42]. A decrease of amino acids is also common in viral infections. During cricket paralysis viral infection, phenylalanine, histidine, isoleucine, valine, proline and leucine, were observed to decrease in concentration in the first 6 hours post-infection, and then levels began increasing after 24 hours, returning to a similar concentration prior to infection after 36 hours [42]. This increase was suggested to be the result of degradation of the cell that viral replication damage has caused, which in turn, resulted in an increase of proteases and hydrolases breaking down proteins and macromolecules [42]. These trends, and decreased levels of pyrimidines and derivatives in DCV-infected insects compared to uninfected insects in this study is a consequence of the increased need of both the host and the virus for adenosine triphosphate (ATP) as the virus continues to utilize available host energy for protein synthesis, and DCV harvesting host nucleotides to replicate viral RNA.

This study also saw an observed decrease of lipids in *w*Mel-free DCV-infected *Drosophila* relative to uninfected *Drosophila* (Fig. 2E). Lipids are one of the key metabolites seen to be modulated by viruses as they utilize them throughout their lifecycle. For example, lipids are often incorporated into the final assembled virus, or in the case of cholesterol, are required for entry, replication, virion assembly, and exit [30, 43–46]. Indeed, Caragata, et al. (2013) determined that DCV-infected Drosophila fed cholesterol-enriched diet resulted in diminished *Wolbachia* pathogen blocking [31]. While the metabolic fate of these lipids cannot be ascertained in this study, it is likely that the lipids decrease in concentration in DCV-infected *Drosophila* because they have been utilized to complete the virus lifecycle. To completely elucidate the metabolic interaction occurring a more focused investigation of the role play in *Wolbachia-*host or DCV-host symbioses lipids is required. Moreover, it would be of interest to compare the findings in this study to the metabolic profiles of insects, including the novel host – *Aedes aegypti*, infected with a variety of other RNA viruses previously proven to confer a protective effect such as dengue-or zika-virus. Given that *w*Mel is the strain employed as a bioagent, it is likely that these same points of conflict are exhibited in the novel *Aedes aegypti* mosquito species. It is this study’s aim that the affected metabolites observed, and the inferences made on how this might contribute to the broad understanding of *Wolbachia*-mediated antiviral protection. Therefore, pinpointing metabolites and metabolic pathways involved in pathogen blocking and support the hypothesis of metabolic interaction as a mechanistic system.

When observing the metabolic profile of DCV infection in *w*Mel-infected *Drosophila* we begin to see evidence of direct metabolic interactions that likely interfere with *Wolbachia*-mediated antiviral protection. The metabolic state of co-infected *Drosophila* at 5 dpi almost mirrors *w*Mel-infected *Drosophila* with no significant difference observed in pairwise comparisons between 5 and 9 dpi *w*Mel-infected *Drosophila* and 5 dpi co-infected *Drosophila* at both (Supplementary Fig. S3 and Table S2). The metabolic state of co-infected *Drosophila* at 9 dpi, however, are different from at both 5 and 9 dpi *w*Mel infected *Drosophila* and likely reflects the beginning of DCV overwhelming the system, where the insect will eventually succumb to the viral infection.

In contrast to single DCV infection, co-infection appeared to drive metabolism to intermediate metabolites resulting in higher levels of glycolytic end products, amino acids, lipids and pyrimidines. This is likely because the metabolites were not immediately utilized by the virus or metabolized to create RNA when compared to single DCV-infected *Drosophila*, as these metabolites are still visible in the system resulting in the increase (Fig. 4). A previous transcriptomics study indicated *w*Mel infected *D. melanogaster* had reduced gene expression of pyrimidine and increased gene expression of purine *de novo* biosynthesis pathways [33]. This could suggest that the reduced gene expression of the pyrimidine pathways is because pyrimidines were being synthesized (or their consumption reduced) in the presence of *w*Mel. The *w*Mel-infected and co-infected *Drosophila* showed an increased abundance of pyrimidines and derivatives would support the down-regulation of those biosynthetic pathways.

The observed increase of nucleotide metabolism in co-infected *Drosophila* suggests that there is likely incomplete protection against DCV-induced death in *w*Mel-infected *Drosophila* when injected, where this metabolic trend would benefit DCV replication. This increase of nucleotides could result in reduced expression of purine salvage pathway, a proposed downstream metabolic consequence that could inhibit virus replication [33]. The incomplete protection is demonstrated in the survival data in this study (Supplementary Fig. 1A) and other studies [6, 7], where the co-infected *Drosophila* eventually succumb to DCV when injected. It is possible that this metabolic profile in *Drosophila* would change to reflect an even more contrasting metabolic profile even more like uninfected *Drosophila,* had the *Drosophila* orally infected with DCV instead of injected. This is because while an injection of DCV ultimately kills the *Drosophila* [6, 7], *w*Mel, however, can clear DCV infection in *Drosophila* when orally infected with DCV [47]. It is also possible as *Wolbachia* consumes host pyrimidines and amino acids (also precursors for nucleotides), the total pool of available metabolites is reduced in turn, slowing the rate at which DCV replicates, and the increase we see is because these metabolites are retained in *Wolbachia*.

By investigating the effect of *w*Mel infection on DCV-infected *D. melanogaster*, this study sought to identify points of metabolic competition between *w*Mel and DCV. When the metabolic states of co-infected *Drosophila* (5 dpi) were compared to both 5 and 9 dpi *w*Mel-infected *Drosophila,* almost no difference was observed; yet DCV infected flies (5 dpi) were significantly different with an inverse metabolic state observed (Fig. 2D). For example, in co-infected *Drosophila* glycolysis was decreased relative to DCV-infected *w*Mel-free *Drosophila,* whereas DCV infection appeared to increase glycolysis relative to uninfected *Drosophila*. The four-way comparison (Fig. 3A) of uninfected, co-infected and singly infected cohorts at 5 dpi, supports the notion that *w*Mel in co-infected *Drosophila* exerts the greatest influence on host metabolism in direct competition with DCV, resulting in a metabolic profile more like *w*Mel-infected *Drosophila* (Fig. 3B) that protects against DCV. This is further supported by *w*Mel’s ability to delay viral accumulation and delayed virus-induced mortality of *Drosophila*, observed in both this study (Fig 1) and previous studies [6, 7].

Time also appeared to affect the metabolic profile in *Drosophila*. While this was not the focus of the differences in the metabolic state of uninfected *Drosophila* at 5 dpi were primarily a hypermetabolic state, where many of the metabolites were increased in concentration relative to uninfected *Drosophila* at 9 dpi (Supplementary Fig. 2B). It is possible that these effects are a result of time. However, it is more likely that this metabolic difference can be attributed to needle-stick injury. Further research beyond this study would be required to examine this effect in more detail.

Taken together the results support that metabolic competition is likely involved in *Wolbachia*- mediated antiviral protection; however, metabolic interaction might be a more apt description of the mechanism behind *Wolbachia*-mediated antiviral protection against RNA viruses. That said, the metabolites identified here might be a point of competition as both microbes drive metabolism in different direction or require these metabolites to complete their life cycles; however, until future targeted studies perhaps manipulating the concentration of these metabolites are conducted definitive statements cannot be made. This is because this study examined the global metabolic profile averaged across entire organisms, for DCV-, *w*Mel-, co-infected and uninfected insects. However, this mechanism does align with previous evidence revealing that *w*Mel’s antiviral protection is a localized response and not systematic [18, 27], and thus is a consequence of *Wolbachia* infection driving host cellular metabolism in competition to DCV. We believe that *w*Mel may drive metabolism in a direction that does not support DCV replication and in doing so, it concomitantly triggers immune pathways via ROS that contribute towards *Wolbachia*-mediated pathogen blocking, and perhaps other downstream metabolic pathways as suggested by Lindsey et al. (2021) [31]. We therefore propose that *Wolbachia*-mediated antiviral protection should be viewed as not just one mechanism but a multimodal response resulting from *w*Mel’s influence on host metabolism.

## Materials and Methods

### Drosophila and Wolbachia

*Two Drosophila melanogaster* fly stocks were maintained on a standard cornmeal diet at a constant temperature of 25°C with a 12-hour light/dark cycle. All experiments used a previously described paired lines of *w*^1118^ flies that were infected by *Wolbachia* (*w*Mel) or had been cured of *Wolbachia* (*w*Mel-free) following a tetracycline treatment (0.3 %) [47]. All experiments were completed after flies had recovered from tetracycline treatment for at least 5-6 generations and gut flora reconstituted [48]. Reverse transcription PCR was used to confirm that all *Drosophila melanogaster* were free of commonly used viruses before experimentation (Supplementary Fig. S5). A standard PCR assay [49] was used to confirm the presence of *Wolbachia* in the *w*Mel line and that all tetracycline treated *D. melanogaster* remained *w*Mel-free (Supplementary Fig. S6).

### Virus

Plaque purified DCV isolate EB was propagated in *w*^1118^ flies and purified by sucrose-gradient purification as described previously [10]. Virus concentration (infectious units per mL; IU/mL) was determined using the 50 % tissue culture infectious dose (TCID50) method [50], as previously described [47]. DCV stocks were diluted in 1× phosphate buffered saline (PBS) to a working concentration of 10^7^ IU/mL, as previously described [11]. DCV stocks were stored at -20°C, and the diluted virus was kept on ice during use to preserve the infectious material.

### Infecting *Drosophila* fly lines with DCV

All DCV infection experiments followed standard protocol as previously described [11]. Briefly, 5- to-8-day old adult male *Drosophila* were anesthetized with carbon dioxide and then injected with 50.6 nL DCV diluted in PBS to a concentration of 1 ×10^7^ IU/mL into the upper lateral part of the abdomen using a Nanoject II microinjector (Drummond Scientific, Pennsylvania, USA). Negative controls were injected with 50.6 nL of PBS. Flies were allowed to recover from injections, and then 10 replicates of 20 adult flies were collected 5 and 9 days post infection (dpi) for all cohorts of flies *w*Mel-free, *w*Mel-infected, DCV-infected, and co-infected. Flies were snap frozen in liquid nitrogen [51] tissue weight recorded for *Drosophila* tissue extraction and processing.

### Survival and viral accumulation assays

To confirm the susceptibility of *w*Mel-free *D. melanogaster* to DCV and the protective effect of *w*Mel, survival and viral accumulation assays were conducted (Supplementary Fig. S1) as previously described [7]. For the survival assay, 20 *Drosophila* from both *w*Mel and *w*Mel-free lines were injected with DCV, with three biological and two technical replicates, including eight negative controls per line. Daily mortality was recorded, censoring deaths within 1 dpi. Survival curves were generated using a Cox mixed-effects model through using R packages (dplyr, coxme, survminer, and survival) [52–56]. The hazard ratio assumption was verified using Cox proportional-hazards regression (coxph) [56]. Following injection with DCV, approximately 50 % survival for *w*Mel-free and *w*Mel-infected *Drosophila* lines were observed at 5 and 9 dpi respectively.

Virus accumulation was estimated using an RT-qPCR assay. Approximately 10-20 *Drosophila* adult males per line were infected with DCV, and 5 *Drosophila* were collected per line at 5 and 9 dpi when applicable. Ten negative control flies were prepared, and the experiment was replicated with three biological and two technical replicates. Prior to DCV RNA quantification, DCV RNA was extracted, and random primed cDNA was synthesized. Samples containing the RpL32 primer mix instead of the DCV primer mix were run in parallel. Controls consisting of no cDNA were also run to test for DNA contamination. DCV RNA was quantified using reverse-transcriptase quantitative PCR with Platinum SYBR Green qPCR SuperMix-UDG, amplifying both DCV (primers; 5’-AGG CTG TGT TTG CGC GAA G-3’ and 5’ -AAT GGC AAG CGC ACA CAA TTA -3’) and RpL32 (primers; 5’ -GAC GCT TCA AGG GAC AGT ATC TG-3’ and 5’-AAA CGC GGT TCT GCA TGA G-3’) genes. The qPCR conditions were 95°C for 2 min, followed by 40 cycles of 95°C for 10 sec, 58°C for 20 sec, and 72°C for 20 sec, with subsequent melt curve analysis. DCV genome copy number was normalized against the *Drosophila* gene *RpL32* using QGene software [57] and a qGene analysis R script [58], setting the amplification cut-off to 1.4 and the standard error cut-off to 25 %. Data normality was tested using the Shapiro-Wilk test from the R package ‘*stats*’ in R studio [52]. Due to non-parametric distribution, a Kruskal-Wallis rank sum test was performed followed by a post-hoc Dunn’s test with Holm-Bonferroni correction using the function ‘kruskal.test’ and ‘*dunn.test*’ from the R package ‘*stats*’ and ‘FSA’ [59] respectively.

Results were graphed using GraphPad Prism [60], allowing for comparison of viral accumulation across collection time points and between *Drosophila* lines.

### *Drosophila* Tissue Extraction and Processing

Before NMR spectral data were acquired, the metabolites in the *Drosophila* tissue were extracted. The acquired samples were homogenized using a TissueLyser II (Qiagen, Venlo, The Netherlands) with three glass beads, and 5 mL of ice-cold solvent of 1:1 acetonitrile:water per gram of tissue for 10 min at 4°C [51]. The samples were centrifuged at 12000 × *g* for 10 min at 4°C, and the supernatant was collected. Extraction was repeated, the two supernatants combined, and then the samples were freeze-dried using an Alpha 3-4LSCbasic freeze-dryer (Christ, Osterode am Harz, Germany). All freeze-dried extracts were stored at -80°C. Freeze-dried extracts were resuspended in 250 μL of NMR buffer: 25 μL of 1.5 M potassium phosphate buffer, pH 7.4; 25 μL of DDD consisting of 1 mM DSS as a chemical shift reference (0 ppm), and 1 mM DFTMP as an internal pH standard in D_2_O; 200 μL of reverse osmosis water [51]. These samples were then centrifuged at 12,000 × *g* for 5 min and 240 μL of each sample was transferred into 3 mL NMR tubes, and 10 μL from each sample were pooled into two quality control samples.

### NMR Data Acquisition

NMR spectra were recorded at 298K on a Bruker Avance III HDX 800 MHz spectrometer equipped with a triple Resonance 5 mm (TCI) Cryoprobe, and a SampleJet automatic sample changer (Bruker, Massachusetts, USA).

One-dimensional (1D) Nuclear Overhauser enhancement spectroscopy (NOESY) spectra were recorded for all samples, as previously described [34]. Using the *noesygppr1d* pulse sequence [RD-90°-*t* -90°-*t*_m_-90°-ACQ] (Bruker Biospin pulse program Library), 4 dummy scans and 256 transients were collected into 65,536 data points at a spectral width of 20.03 ppm. With a mixing time (t_m_) of 0.01 sec and relaxation delay of 4 sec. The radiofrequency offset value was set to the frequency of the water resonance.

To aid in metabolite identification, two-dimensional (2D) spectra were also acquired on a pooled quality control sample including: ^13^C-heteronuclear single quantum coherence (HSQC); ^1^H-^1^H-total correlation spectroscopy (TOCSY); ^13^C-HSQC-TOCSY, and ^13^C-heteronuclear multiple bond correlation (HMBC) spectroscopy. The heteronuclear 2D spectra were acquired with the ^13^C carrier frequency set to 110 ppm, and GARP decoupling of the ^13^C channel during the acquisition time. The ‘Echo-Antiecho’ acquisition mode was used in the HSQC and HSQC-TOCSY experiments, and the mode ‘States-TPPI’ was used for the HMBC and TOCSY experiments. The homonuclear experiments, barring the TOCSY, were acquired with 96 transients, 32 dummy scans, and 1.5 sec relaxation delay. The TOCSY and the HMBC had relaxation delays of 2 sec and 1.5 sec; 16 and 256 transients, and 16 and 32 dummy scans, respectively. The mixing time for the TOCSY and HSQC-TOCSY was both 0.08 sec. The data points for the HSQC, TOCSY, HSQC-TOCSY and HMBC was 2,048 × 3000; 4,096 × 400; 4,096 × 300, and 2,048 × 256, respectively. For all the experiments the spectral width was 13.00 ppm for ^1^H NMR and ∼300 ppm for ^13^C NMR.

1D spectra were processed using TopSpin [61], as previously described [34]. The spectra were zero filled twice and the Free Induction Decay (FID) was multiplied by an exponential window function with 0.3 Hz line broadening before Fourier transformation. The acquired spectra were manually phased and baseline corrected and calibrated to the DSS signal at 0 ppm in TopSpin [61]. The resulting NMR signals were corrected for pH and ionic strength-dependent shift variation using the *Icoshift* algorithm in MATLAB [62]. 2D NMR spectra were processed by two-dimensional Fourier transformation after the indirect dimension was multiplied by a squared sine bell window function shifted *π*/2. A Lorentz to Gauss transformation with a line broadening factor of -10 Hz and Gaussian broadening factor of 0.1 was applied in the direct dimension. Phase and baseline correction were applied manually.

### Multivariate Statistical Analysis

The ^1^H NMR spectra were bucketed into 0.001 ppm regions using an in-house bucketing script in MATLAB, across a range of 10.00 to 0.25 ppm. The chemical shift region between 5.10-4.70 ppm was excluded to eliminate any effects of imperfect water suppression. The bucketed data matrices were normalized to the total intensity and imported into the SIMCA 18 package [63] for multivariate statistical analysis.

The data matrix with the bucketed 1D spectra was analyzed using unsupervised principal components analyses (PCA) with Pareto scaling to determine if there were any outliers among the samples and to first investigate any clustering of the cohorts. Partial Least Squares (PLS) with Pareto scaling was then used to maximize the distinction between the cohorts. Meta-data was also provided during these analyses. Each cohort was assigned a number identity and designations where pooled quality controls; co-infected at 5 dpi (+W+V@5); DCV-infected at 5 dpi (-W+V@5); *w*Mel-infected at 5 dpi (+W-V@5); uninfected at 5 dpi (-W-V@5); co-infected at 9 dpi (+W+V@9); *w*Mel-infected at 9 dpi (+W-V@9), and uninfected at 9 dpi (-W-V@9) were 0, 1, 2, 3, 4, 5, 6 and 7 respectively. PCA and PLS model quality was determined through the parameters: R^2^X, R^2^Y, and Q^2^. Where R^2^X and R^2^Y represent the variance in X and Y explained by the model’s latent components, while Q^2^, indicates predictive ability. All PLS models were further validated by 200 random permutation tests, and cross-validated ANOVA (Supplementary Table S1 and Fig. S7).

PCA models were used to characterize the global differences of *w*Mel infection, DCV infection, and co-infections (*w*Mel and DCV), in which multiple models were fitted (**M1-M2**). PCA models were also performed for all cohort at 5 dpi (**M14**), and all *w*Mel (**M15**). Pairwise PCA models compared the effect of age on the metabolic profiles of the *Drosophila* cohorts (PCA: **M3-M5**); the effect of *Wolbachia* infection (PCA: **M6-M7**); the effect of DCV infection (PCA: **M8, M10, M11**), and finally the effect of co-infection (PCA: **M9, M12, M13**) (Supplementary Fig. S8). These comparisons were chosen to best investigate the changes in metabolic profiles between cohorts.

PLS models were used for the pairwise comparisons so that each model could effectively be transformed into bivariate 1D loadings plots using an in-house MATLAB script. Where the loadings coefficients (*p*) back-scaled with the square root of the standard deviation of the variables were plotted against the chemical shift values of the spectral data as previously described [64]. The absolute values of the correlation-scaled loadings coefficients |*p*(*corr*)| were then incorporated into this plot as a heatmap color scale. Variables that were significantly altered in a model were identified when, |*p*|≥ 0.015; |*p*(*corr*)| ≥ 0.6), and VIP ≥ 1. The metabolites were scored for variable importance of projection (VIP), to determine statistically significant metabolites involved in the biological interactions [65].

### Metabolite Identification

The metabolite assignment (Supplementary Table S6) of the NMR peaks was performed using Chenomx NMR Suite [66], with the aid of the 2D spectra, the Human Metabolome Database (HMBD) [67], and the Biological Magnetic Resonance Data Bank (BMRB) [68]. Assignment of metabolites to peaks was also aided by calculating the covariance matrix (statistical total correlation spectroscopy) over the entire spectral region for all NMR spectra using an ‘in-house’ MATLAB script, and overlaying reference 2D spectra from BMRB in the software from the collaborative computation project for NMR (CCPN) [69].

### Univariate Statistical Analysis

To confirm that levels of the metabolites identified in the bivariate loadings plots (|*p*|≥ 0.015; |*p*(*corr*)| ≥ 0.6), and VIP ≥ 1) were either higher or lower in one cohort relative to another cohort, a univariate analysis was also performed as previously described [34]. Only ^1^H NMR peaks from metabolites were selected that had minimal spectral overlap (Supplementary Table S7). Using an in-house MATLAB script, the area integration of the metabolite peaks was found for each non-normalized full resolution spectrum. The data for each metabolite was then grouped by the meta-data with the averages calculated ± the standard deviation (Supplementary Table S8 and Table S9). Area integration values below a limit of quantification (LOQ) were excluded. The LOQ was determined by calculating the number of data points within integration regions, multiplying this by the average noise level to obtain the total error, and then multiplying the total error by 5 (where the noise level contributes more than 20 % error to the area integration) [34]. The average noise level was calculated as the standard deviation of baseline noise between 10-9.8 ppm for each sample spectrum which was then averaged across spectra.

Using the R packages ‘*ggplot2*’ [70], ‘*car*’ [71], ‘*dunn.test*’ [72], and ‘*ggsignif*’ [73], in an in-house Rscript in R studio [52], a Kruskal-Wallis statistical analysis was performed on the integration means followed by a Dunn post-hoc test with Holm-Bonferroni correction comparing each of the cohorts for each metabolite plotted as a violin plots for each metabolite (Supplementary Fig. S3 and Table S2).

## Supporting information

Supplementary Information: supplementary text, figures S1 to S8, tables S1 to S9, and SI references

## Acknowledgments

We thank Dr. Angelique Asselin and Dr. Daniel Chew for their assistance with completing the survival experiments. We thank Ms. Alexandra Gloria, Mr. Luke Husdell, and Mr. Kern Webster for their advice and comments on the manuscript.

## Supplementary Material

Supplementary material available at [Bioarchive] online.

## Data Availability

The NMR data presented in this study can be found in the online repository – MetaboLights. It can be found under the accession number ####.

## Author Contributions

Conceived the experiments: K.N.J, J.C.B. and H.J.S. Conducted the insect injection experiments: S.M.W. and T.G.T. NMR experiments were completed by S.M.W. Data analysis, interpretation and preparation of the manuscript: S.M.W, T.G.T, K.N.J, J.C.B., and H.J.S.

**Competing Interest Statement**

The authors declare no competing interests.

