## Supplementary Information: supplementary text, figures S1 to S8, tables S1 to S9, and SI references for "*Wolbachia*-mediated antiviral protection is driven by its multimodal effects on *Drosophila melanogaster* metabolism"

### Supplementary Information for
*Wolbachia*-mediated antiviral protection is driven by its multimodal effects on *Drosophila melanogaster* metabolism

Sarah M. Walsh[1,2], Tyson G. Thomson[3], Karyn N. Johnson[3], Jeremy C. Brownlie[1], Horst Joachim Schirra[1,2,4]*

[1] School of Environment and Science, Griffith University, Brisbane, QLD, Australia
[2] Institute for Biomedicine and Glycomics, Griffith University, Brisbane, QLD, Australia
[3] School of the Environment, The University of Queensland, Brisbane, QLD, Australia
[4] Centre for Advanced Imaging, The University of Queensland, Brisbane, QLD, Australia

*Horst Joachim Schirra.

**This PDF file includes:**

Supplementary text
Figures S1 to S8
Tables S1 to S9
SI References

**Supplementary Information Text**

**Confirmation of the effect of a Drosophila C virus in *D. melanogaster***

To confirm approximate days of 50 % mortality for *w*Mel and *w*Mel-free Fly lines infected with Drosophila C virus (DCV) a survival assay was performed. This assay also reconfirmed *w*Mel reduced DCV pathology in the fly lines used. The survival assay confirmed that when injected *D. melanogaster* were susceptible to DCV (HR = 5.42, P<0.001 at p = $1.3 \times 10^{-15}$, Cox mixed-effects model; Supplementary Fig. S1A), where the assumption that the hazard ratio remains the same overtime was determined to be valid ($\chi^2(1, N = 204) = 0.72$, p = 0.4) where p > 0.05 (Cox proportional-hazards regression model). *w*Mel increased survival rates against DCV, where in *w*Mel-free *Drosophila* DCV induced 100% mortality within 9 days post infection (dpi), while *w*Mel-infected *Drosophila* survived until day 19 dpi (HR = -1.89, P<0.001 at p = 0, Cox mixed-effects model; Supplementary Fig. S1A). Approximately 50 % survival for *w*Mel-free and *w*Mel-infected *Drosophila* lines were observed at 5 and 9 dpi respectively (Supplementary Fig. S1A). The time points 5 and 9 days post infection, were used for all the proceeding experiments, including the viral accumulation assay (Supplementary Fig. S1B).

The delay in DCV-induced mortality corresponded to a delay in viral accumulation, where *w*Mel delays DCV accumulation, as well as delays virus-induced death (Supplementary Fig. S1B). There was significant difference of DCV accumulation for *Drosophila* infected with *w*Mel and *w*Mel-free *Drosophila* ($H(2) = 10.288$, p = 0.0058, 95% CI [0.65, 1.00], $n_{obs} = 16$; Supplementary Fig. S1B). The mean normalized expression of DCV accumulation was statistically different in two of the three pairwise comparisons. There was a significant difference ($p < 0.05$) in viral RNA accumulation between *w*Mel-free *Drosophila* and *w*Mel-infected *Drosophila* collected at 5 dpi (p = 0.005) (Supplementary Fig. S1B). There was no statistical difference ($p < 0.05$) in DCV accumulation between 5 and 9 dpi *w*Mel-infected flies (Supplementary Fig. S1B).

**Virus-free Confirmation**

Reverse transcriptase PCR (RT-PCR) was used to confirm the absence of contaminating virus in the flies before experimentation, as previously described [1]. For this assay, a pool of six flies from each *Drosophila* line (*w*Mel and *w*Mel-free) was collected. These samples were lysed and homogenized as above, but using the TRIzol reagent (Thermo Fisher Scientific, Massachusetts, USA) instead of STE, to separate out the phase containing RNA. Following phase separation and the isolation of the RNA, the nucleic acid was precipitated by isopropanol and then washed to remove any remaining reagents using 70% ethanol. Total RNA was resuspended in RNase-free water. RNA purity and concentration were determined using the Epoch Microplate Spectrophotometer (Biotek, Vermont, USA).

Samples were DNase treated to avoid genomic DNA contamination. For this, the samples were first diluted to 2 µg total RNA in 5.75 µL total volume. The extracted RNA was treated with 2 µL RQ1 RNase-free DNase, 1 µL RQ1 DNase 10× Reaction Buffer, and then 1 µL RQ1 DNase Stop Solution (Promega, Wisconsin, USA) for each reaction to inactivate the enzyme.

Reverse transcriptase was used to synthesize complementary DNA (cDNA) of the RNA samples for PCR. This was performed using SuperScript III Reverse Transcriptase (Invitrogen, Thermo Fisher Scientific, Massachusetts, USA), with 0.25 µL random primers [250 µM] and 0.5 µL dNTPs [10 µM]. The priming incubation occurred for 5 min at 60 °C and at 4 °C for 2 min. Reverse transcription was then initiated by adding 2 µL 5× First-Strand Buffer; 0.75 µL RNase-free water; 0.5 µL Dithiothreitol, and 0.25 µL SuperScript III Reverse Transcriptase (RT), to each sample.

PCR of the cDNA was then conducted following a similar procedure as described for 'Confirmation of *w*Mel infection status'. However, the primers used were for DCV, Flock House virus, Cricket Paralysis virus and Sindbis virus, as previously described [2] (Supplementary Table S3). A separate cDNA sample was also run with the RpL32 primers to confirm the presence of cDNA. Positive and negative controls consisting of cDNA previously synthesized from *Drosophila* were injected with the viruses (positive controls), and non-template controls (negative controls) were also run. This assay was visualized as per 'Confirmation of *w*Mel infection status', identified by the expected amplicons (Supplementary Fig. S5).

**Confirmation of *w*Mel infection status**

The absence or presence of *w*Mel was confirmed using a standard PCR assay [3].Therefore, five adult mixed-sex flies were collected from each of the *w*Mel-free and *w*Mel *Drosophila* lines. These pool of five flies were homogenized in 50 µL of sterile STE buffer [100 mM NaCl; 10 mM Tris Cl, pH 8.0; 1 mM EDTA, pH 8.0], using the STE method of DNA extraction previously described [3], and a TissueLyser II (Qiagen, Venlo, The Netherlands) with three glass beads at 30 Hz for 10 sec. This procedure was repeated with an STE blank control. All the samples were supplemented with 2 µL of proteinase K [10 mg/mL] and centrifuged at

14000 × *g* for 2 min. All samples were transferred to PCR tubes and incubated for 30 min at 37 °C and again for 10 min at 95 °C. All samples were kept on ice until the PCR was performed.

The PCR assay was conducted using 23 µL of master mix [5 µL of myTaq buffer (Meridian Bioscience, Ohio, USA); 0.5 µL of 10 µM *wsp* primer mix or RpL32 primer mix (Supplementary Table S4); 0.25 µL of myTaq Polymerase, and 16.75 µL of sterile water]. A PCR temperature profile of 94°C for 30 sec, 55°C for 30 sec, and 72°C for 1 min was used for 35 cycles. Various controls were also assayed simultaneously, as outlined in Supplementary Table S5.

The PCR products were visualized employing electrophoresis and using a Mini-Sub Cell GT Cell (BioRad, California, USA). 1.3% agarose gel stained with SYBR Safe DNA Gel Stain (Invitrogen, Thermo Fisher Scientific, Massachusetts, USA) was used and a 100 bp DNA ladder (Meridian Bioscience, Ohio, USA) was utilized as a reference. Loading was aided by Purple Gel Loading Dye (NEB, Victoria, Australia). The expected amplicons of the PCR products for RpL32 and *wsp*, 143 and 181 bp respectively, were used to confirm absence or presence of *w*Mel (Supplementary Fig. S6).

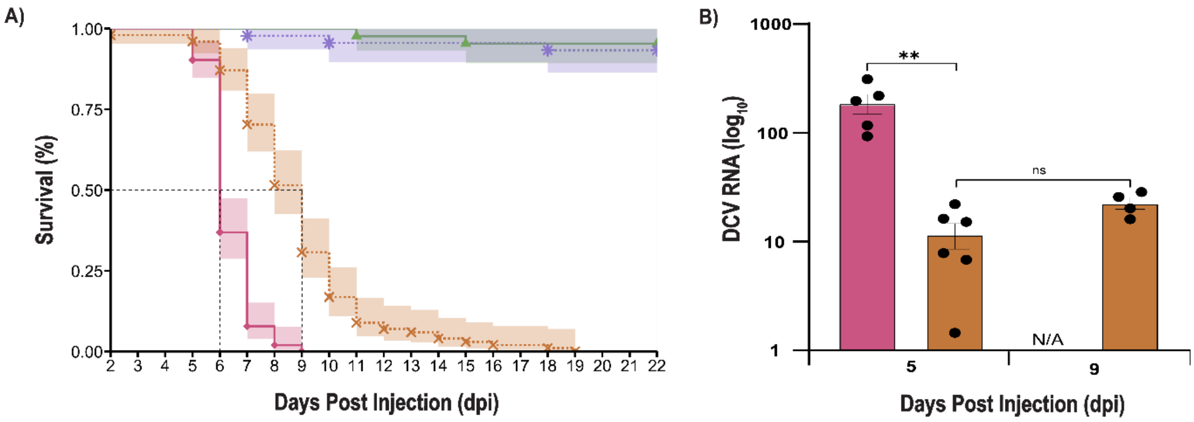


**Supplementary Figure S1:** *Wolbachia* strain *w*Mel provides antiviral protection in *D. melanogaster*. A) Survival of DCV-infected (pink and orange) and PBS-injected (purple; asterix, and green; triangle) *Drosophila* with (broken lines) or without (solid lines) *w*Mel infection, where *w*Mel-infected (orange; cross) *Drosophila* showed significantly extended survival (P<0.001) compared to *w*Mel-free (pink; diamond) *Drosophila* when infected with DCV. (shaded areas: 95 % confidence intervals). B) Mean viral accumulation of DCV in *w*Mel-infected (orange bar) and *w*Mel-free (pink bar) cohorts infected with DCV collected at 5 and 9 dpi at a logarithmic scale ($log_{10}$) relative to *RpL32*. Column bars represent the mean with SEM shown; replicate data indicated by black dots. Due to high mortality by 9 dpi, *w*Mel-free *Drosophila* infected with DCV were not collected for analysis, indicated as N/A. The outcome of statistical analysis is indicated with P values: α = 0.01 is '**', and non-significant values are 'ns'.

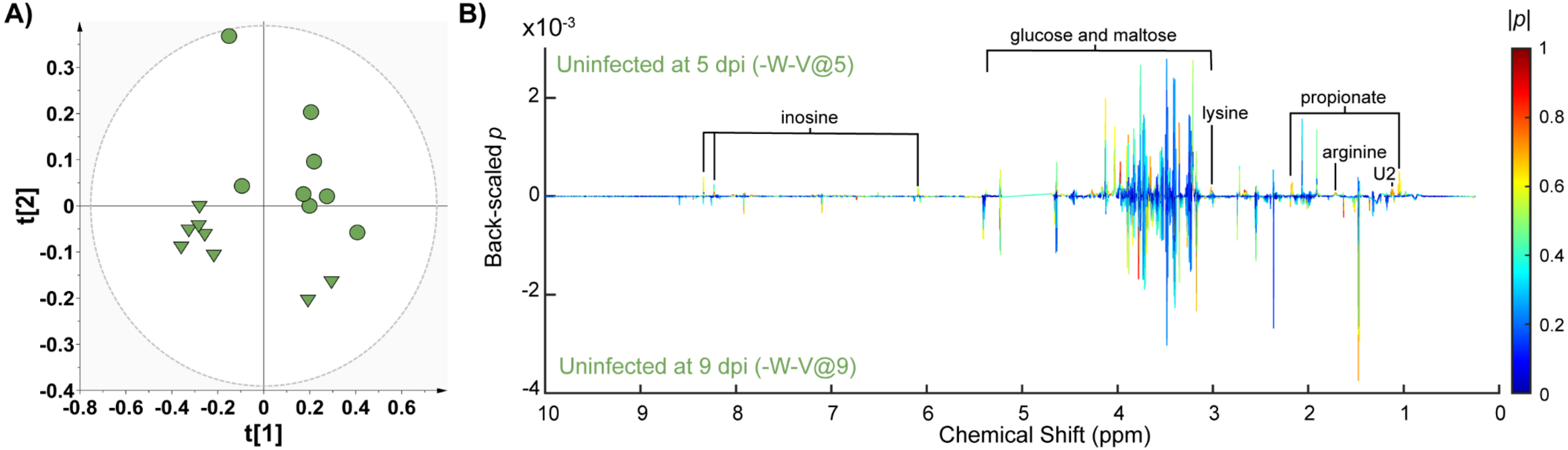


**Supplementary Figure S2:** Age comparison for uninfected *Drosophila*. PLS scores and bivariate loadings plots of uninfected *Drosophila* at 5 dpi compared to uninfected *Drosophila* at 9 dpi. A) Scores plots, where component 1 on the x-axis is indicated by t[1] and component 2 is on the y-axis is t[2]. B) Bivariate loadings plots, where the back-scaled loadings coefficients *p* was plotted against the chemical shift (ppm) for the bucketed multivariate statistical analysis *X*-matrices' variables obtained from the 1D $^{1}$H NMR spectra. The correlation coefficients |*p*(*corr*)| were superimposed on the bivariate loadings plot as a heatmap color scale. This figure follows the same color and shape scheme as previous multivariate statistical analysis models. Metabolites that were significantly altered between comparisons in the univariate analysis were annotated in black ($p \leq 0.05$).

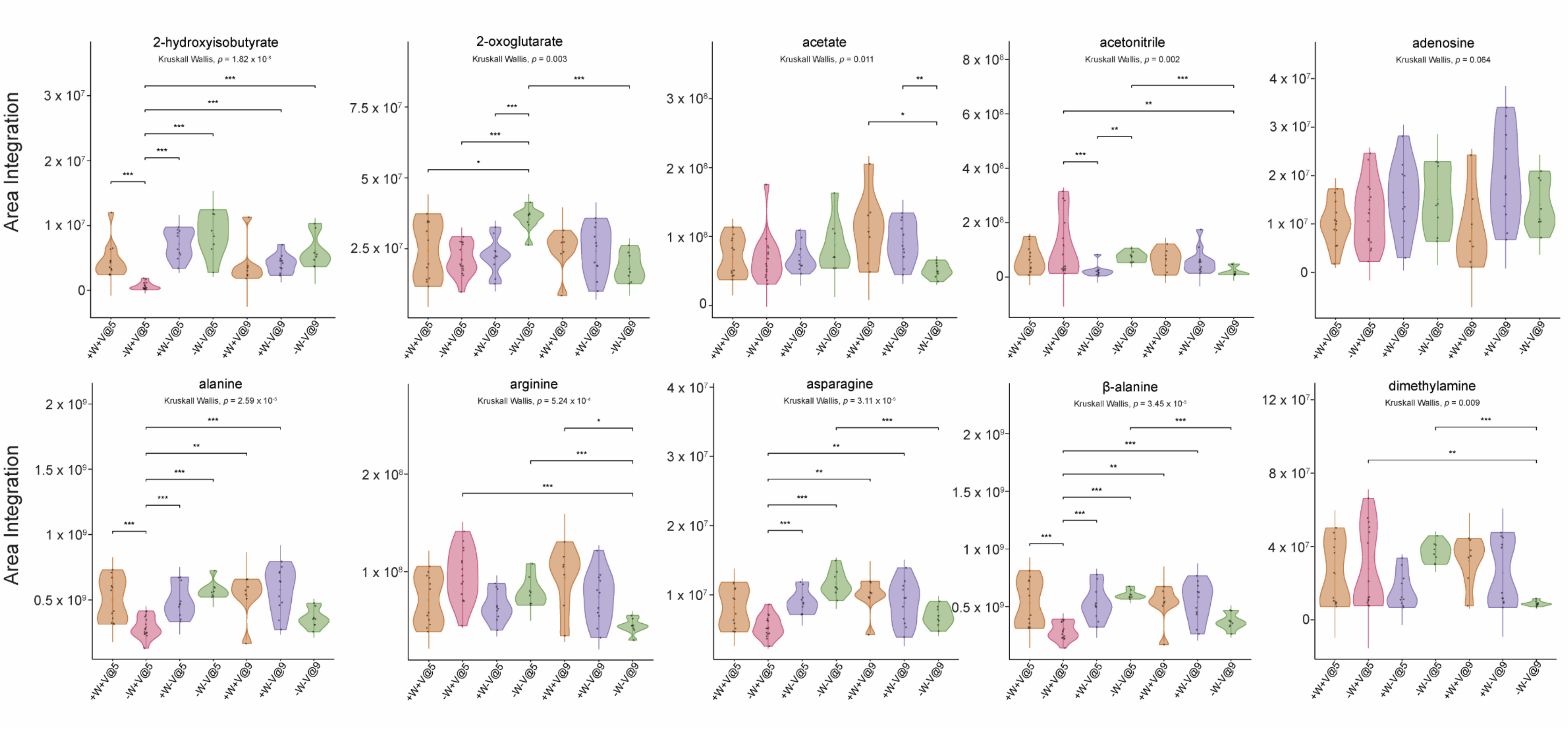


**Supplementary Figure S3A:** Violin Plots of Concentration Changes in Metabolites Part A. Where the x-axis is the meta-Y-table identity of the cohorts. As illustrated groups that were significantly significant different, as determined by the post-hoc Dunn test with Holm-Bonferroni correction, is displayed with common significant codes: $p < 0.05$ is ‘*’, $p < 0.01$ is ‘**’, and $p < 0.001$ is ‘***’.

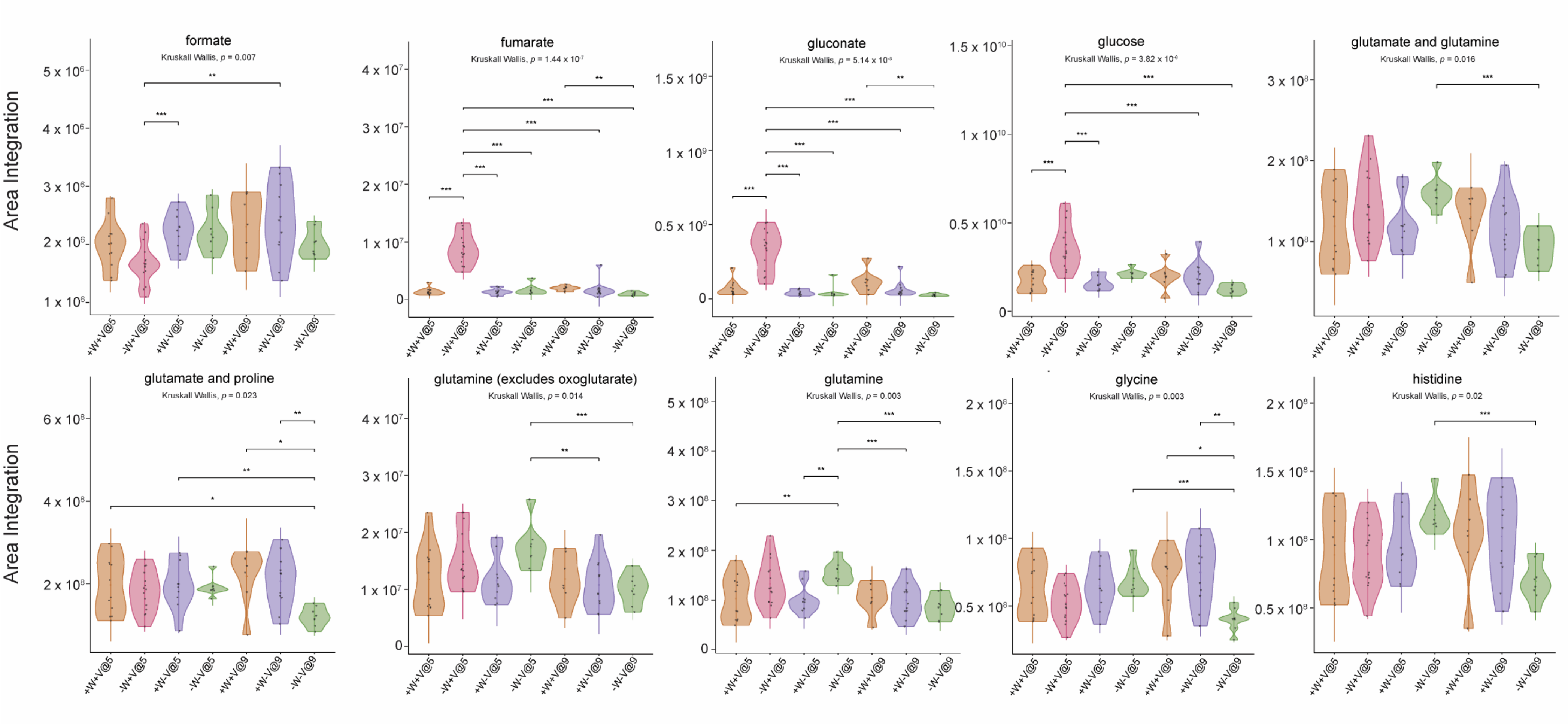


**Supplementary Figure S3B:** Violin Plots of Concentration Changes in Metabolites Part B. Where the x-axis is the meta-Y-table identity of the cohorts. As illustrated groups that were significantly significant different, as determined by the post-hoc Dunn test with Holm-Bonferroni correction, is displayed with common significant codes: p < 0.05 is ‘*’, p < 0.01 is ‘**’, and p < 0.001 is ‘***’.

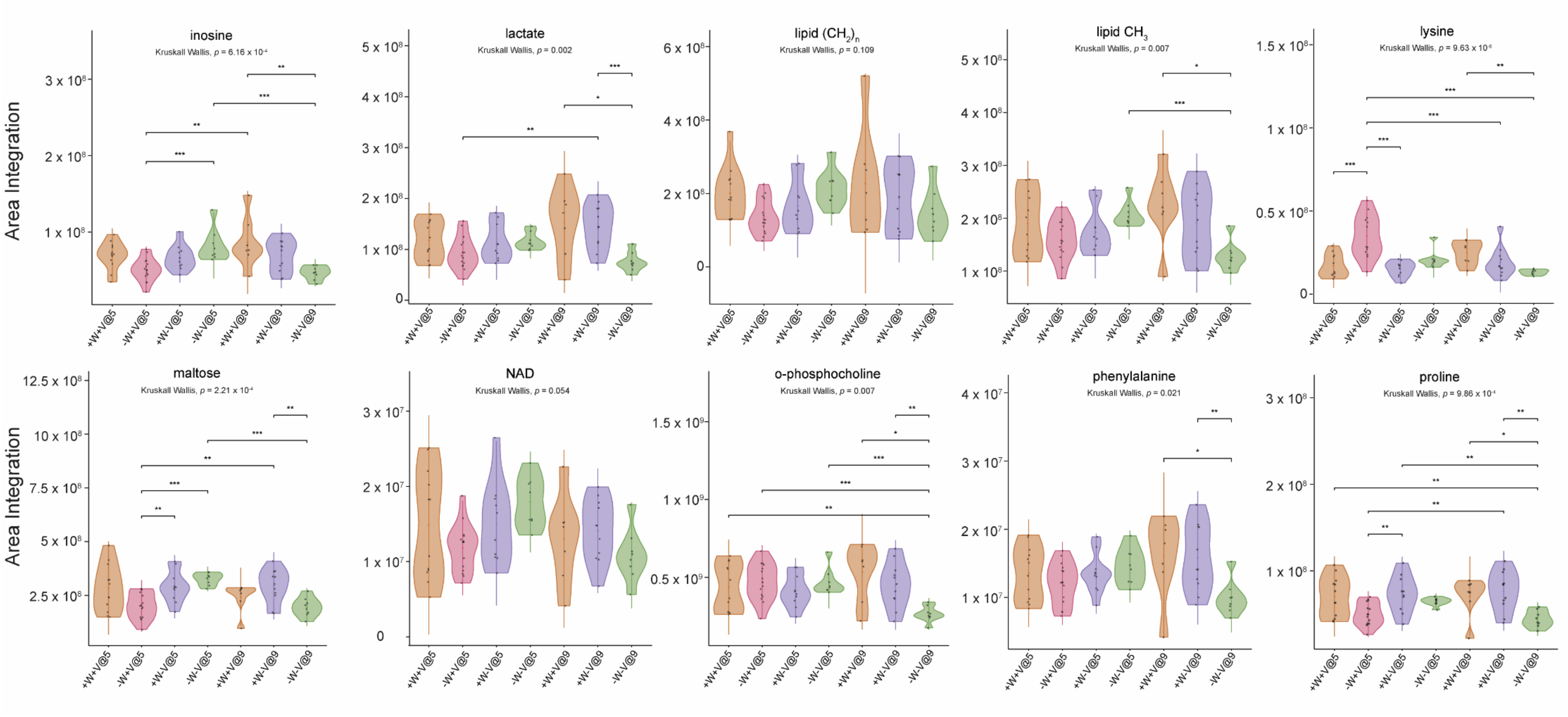


**Supplementary Figure S3C:** Violin Plots of Concentration Changes in Metabolites Part C. Where the x-axis is the meta-Y-table identity of the cohorts. As illustrated groups that were significantly significant different, as determined by the post-hoc Dunn test with Holm-Bonferroni correction, is displayed with common significant codes: $p < 0.05$ is '*', $p < 0.01$ is '**', and $p < 0.001$ is '***'.

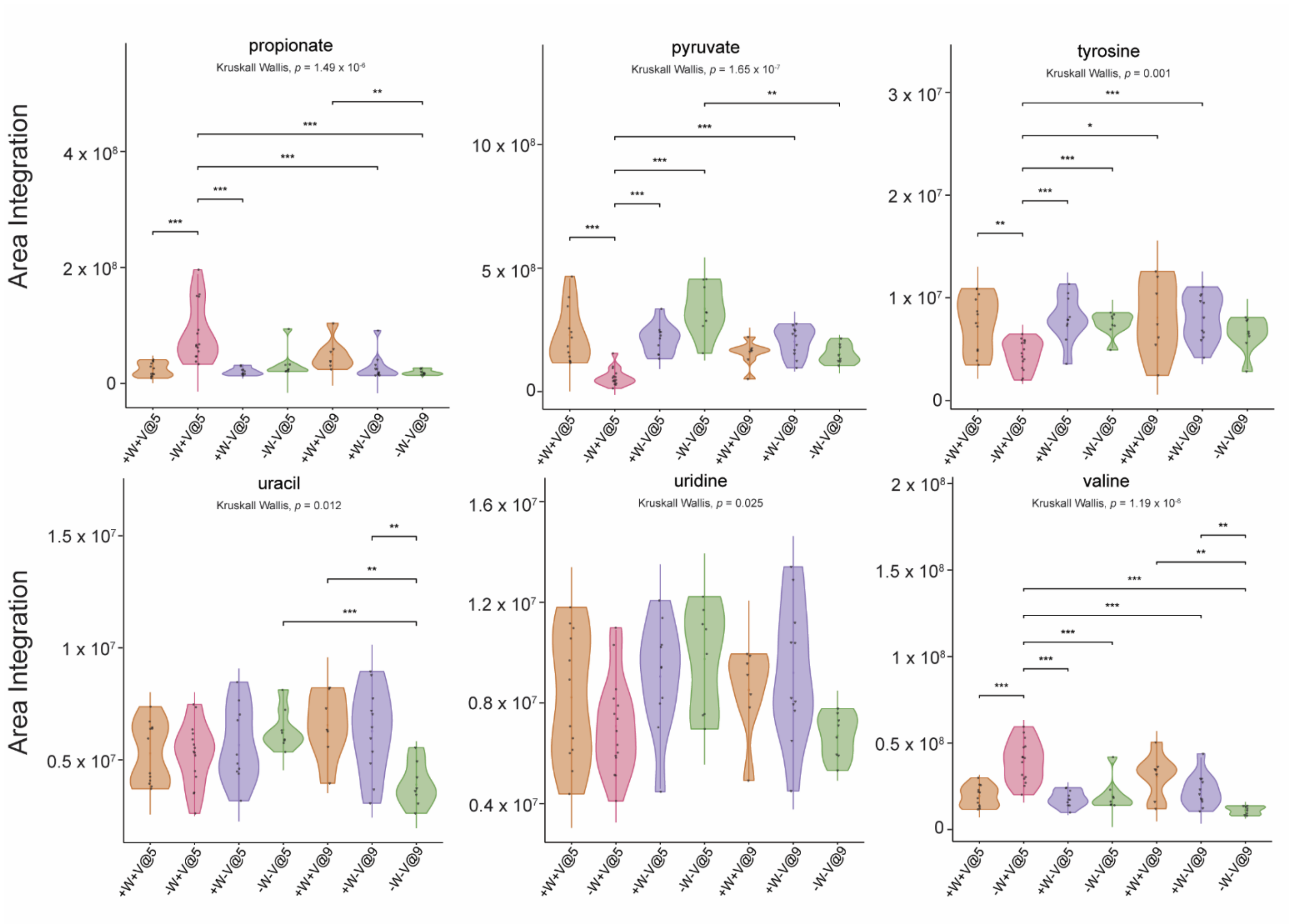


**Supplementary Figure S3D:** Violin Plots of Concentration Changes in Metabolites Part D. Where the x-axis is the meta-Y-table identity of the cohorts. As illustrated groups that were significantly significant different, as determined by the post-hoc Dunn test with Holm-Bonferroni correction, is displayed with common significant codes: $p < 0.05$ is '*', $p < 0.01$ is '**', and $p < 0.001$ is '***'.

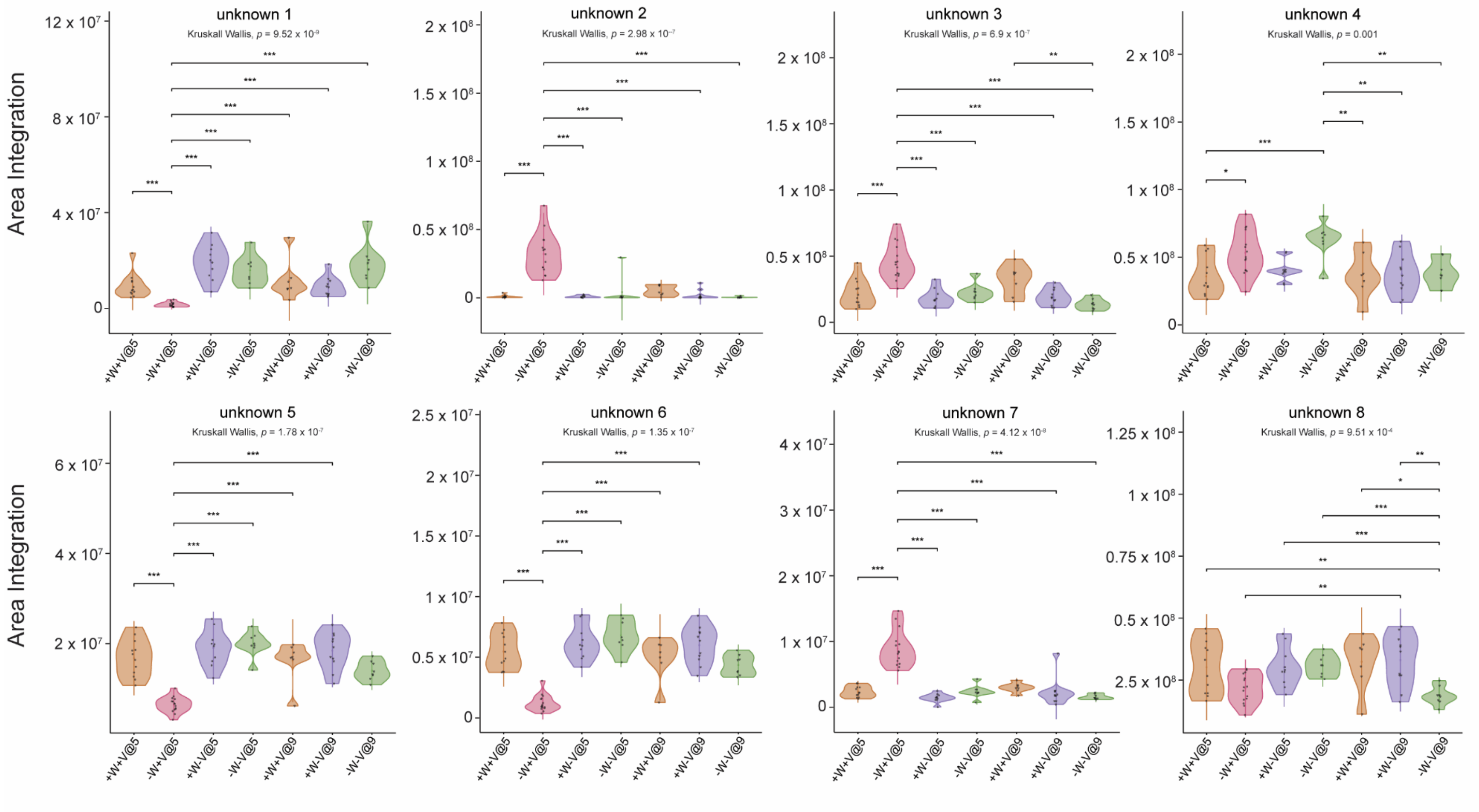


**Supplementary Figure S3E:** Violin Plots of Concentration Changes in Metabolites Part E. Where the x-axis is the meta-Y-table identity of the cohorts. As illustrated groups that were significantly significant different, as determined by the post-hoc Dunn test with Holm-Bonferroni correction, is displayed with common significant codes: $p < 0.05$ is ‘*’, $p < 0.01$ is ‘**’, and $p < 0.001$ is ‘***’

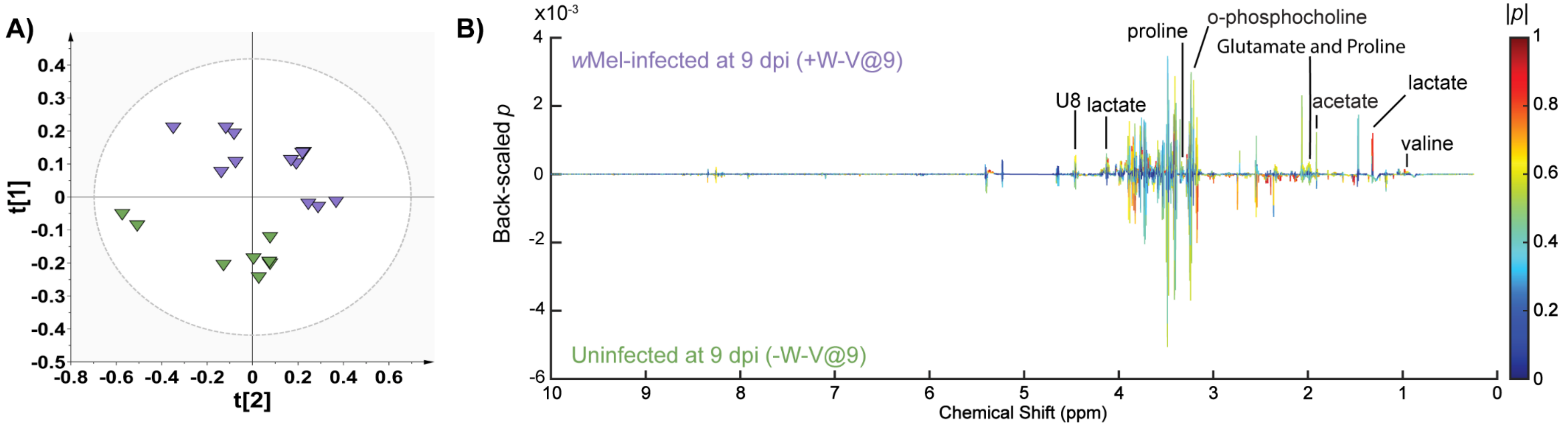


**Supplementary Figure S4:** PLS scores and bivariate loadings plots of *Wolbachia*-infected *Drosophila* at 9 dpi compared to uninfected *Drosophila* at 9 dpi. A) Scores plots, where component 1 on the x-axis is indicated by t[1] and component 2 is on the y-axis is t[2]. B) Bivariate loadings plots, where the back-scaled loadings coefficients *p* was plotted against the chemical shift (ppm) for the bucketed multivariate statistical analysis *X*-matrices' variables obtained from the 1D $^{1}H$ NMR spectra. The correlation coefficients |*p*(*corr*)| were superimposed on the bivariate loadings plot as a heatmap color scale. This figure follows the same color and shape scheme as previous multivariate statistical analysis models. Metabolites that were significantly altered between comparisons in the univariate analysis were annotated in black ($p \leq 0.05$).

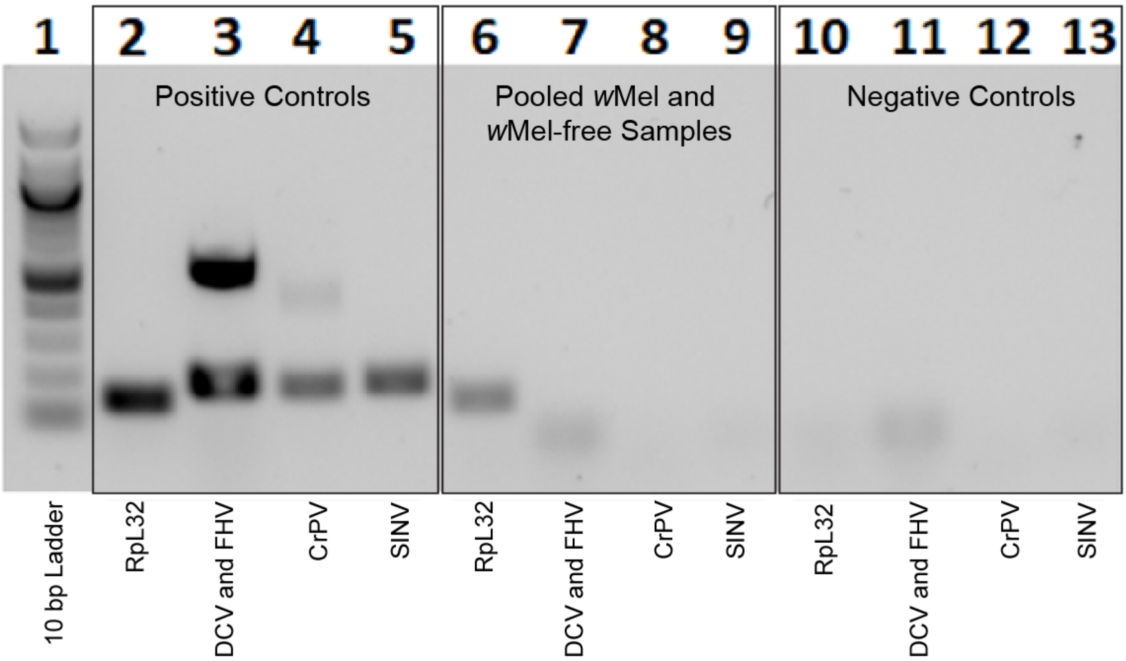


**Supplementary Figure S5:** Virus-free confirmation of *Drosophila* used in study. Both *Drosophila* lines used in this study were determined to be virus-free before experimentation. Common contaminating viruses were screened for in each line (*w*Mel and *w*Mel-free). As indicated the well numbered 1 contains the 100 bp Ladder; wells 2-5 are the positive controls; wells 6-9 are screened lines, and wells 10-13 are the negative controls. The common contaminating viruses include Drosophila C virus (DCV), Flock House virus (FHV); Cricket paralysis virus (CrPV); Sindbis virus (SINV). As visualized in the figure, there was no amplification observed in the pooled *Drosophila* lines and thus confirming the absence of contaminating viruses.

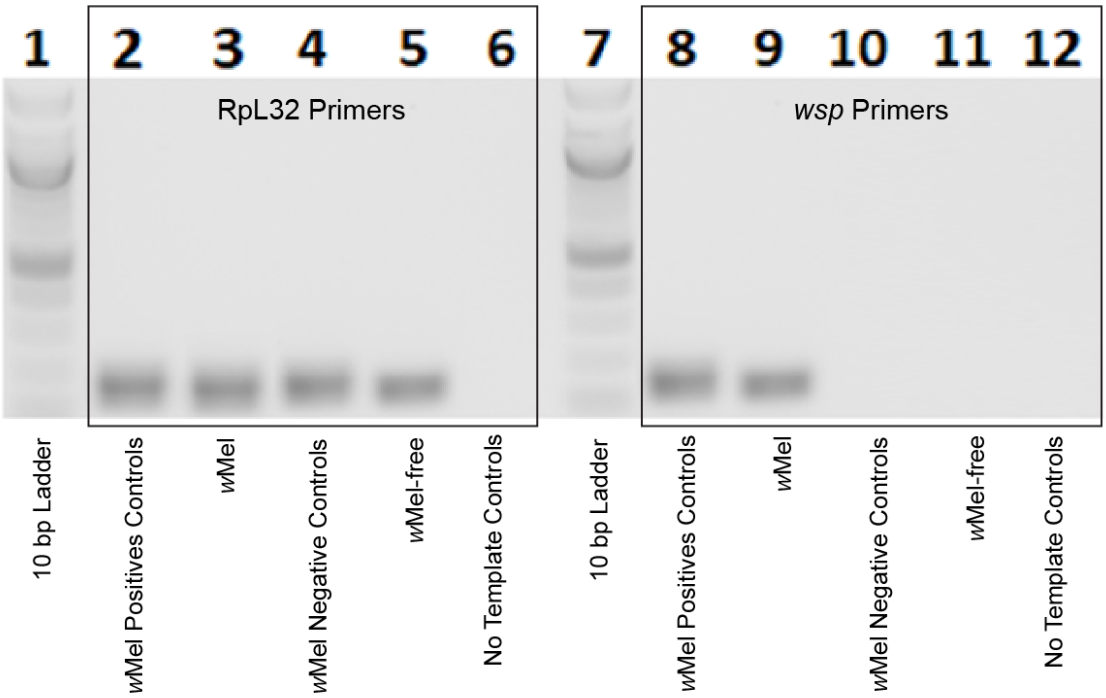


**Supplementary Figure S6:** The absence of *w*Mel was confirmed in the maintained *w*Mel-free *Drosophila* line. Each *Drosophila* line (*w*Mel and *w*Mel-free) was screened for the appropriate presence or absence of *w*Mel. The presence of *w*Mel was confirmed in the *w*Mel-infected line and the absence of *w*Mel was confirmed in the *w*Mel-free line. As indicated the well numbered 1 and 7 contain 100 bp Ladders; wells 2 and 8 contain *w*Mel positive controls; wells 3 and 9 contain the *w*Mel-infected *Drosophila* samples; wells 4 and 10 contain *w*Mel negative controls; wells 5 and 11 contain the *w*Mel-free *Drosophila* samples, and wells 6 and 12 contain the no template controls. Wells 2-6 confirmed the presence of *D. melanogaster* genomic DNA in both *Drosophila* lines using the RpL32 primers. Wells 8-12 confirmed the absence or presence of *w*Mel using *wsp* primers. Well 9 confirms the presence of *w*Mel in the *Drosophila* line and well 11 confirms the absence of *w*Mel in the *w*Mel-free line.

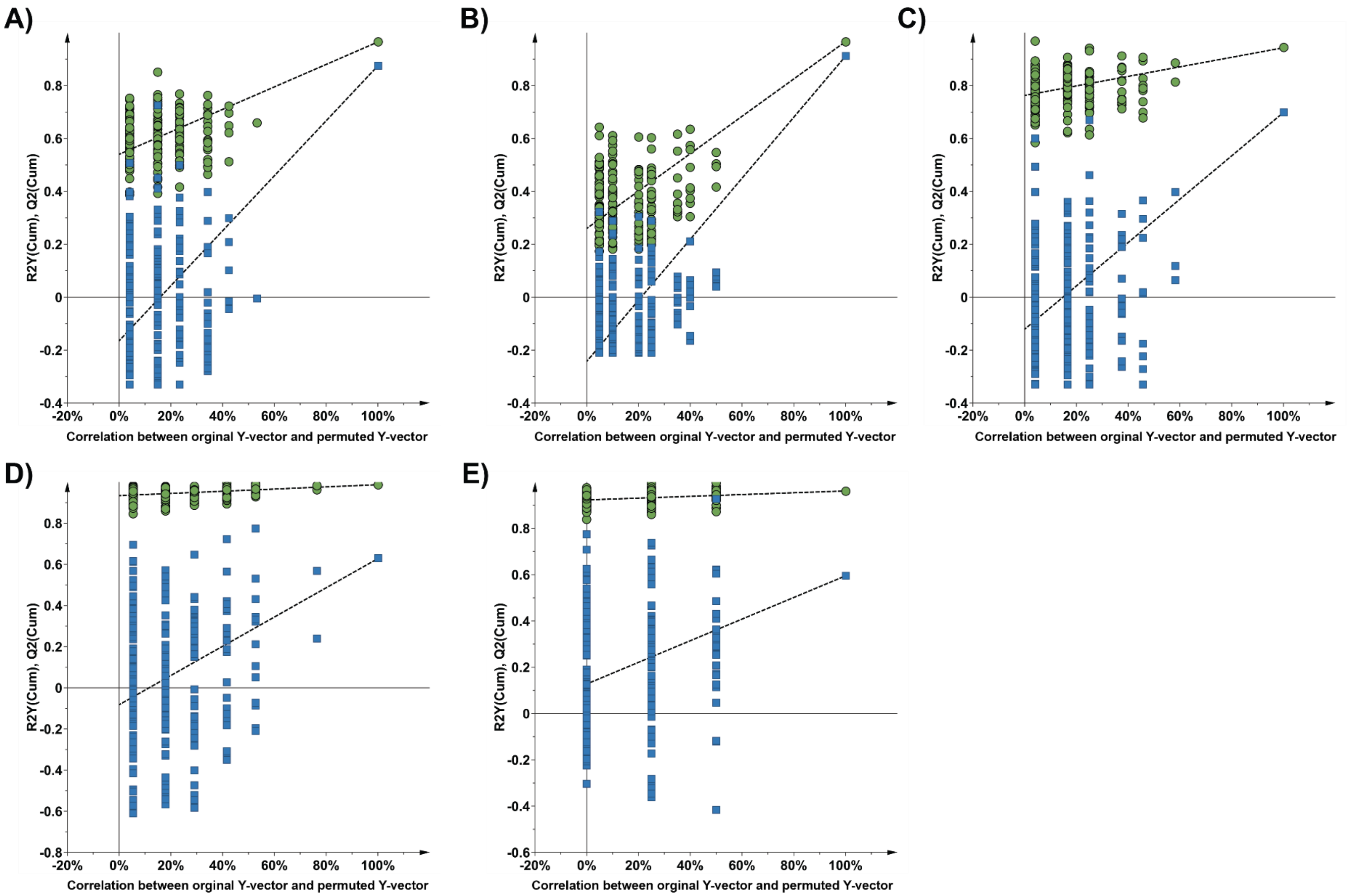


**Supplementary Figure S7:** Permutation Plots A) Uninfected *Drosophila* at 5 dpi and DCV-infected *Drosophila* at 5 dpi. B) Co-infected *Drosophila* at 5 dpi and DCV-infected *Drosophila* at 5 dpi. C) *w*Mel-infected *Drosophila* at 9 dpi and uninfected *Drosophila* at 9 dpi. D) *w*Mel-infected *Drosophila* at 5 dpi and uninfected *Drosophila* at 5 dpi. E) Uninfected *Drosophila* at 5 dpi and uninfected *Drosophila* at 9 dpi.

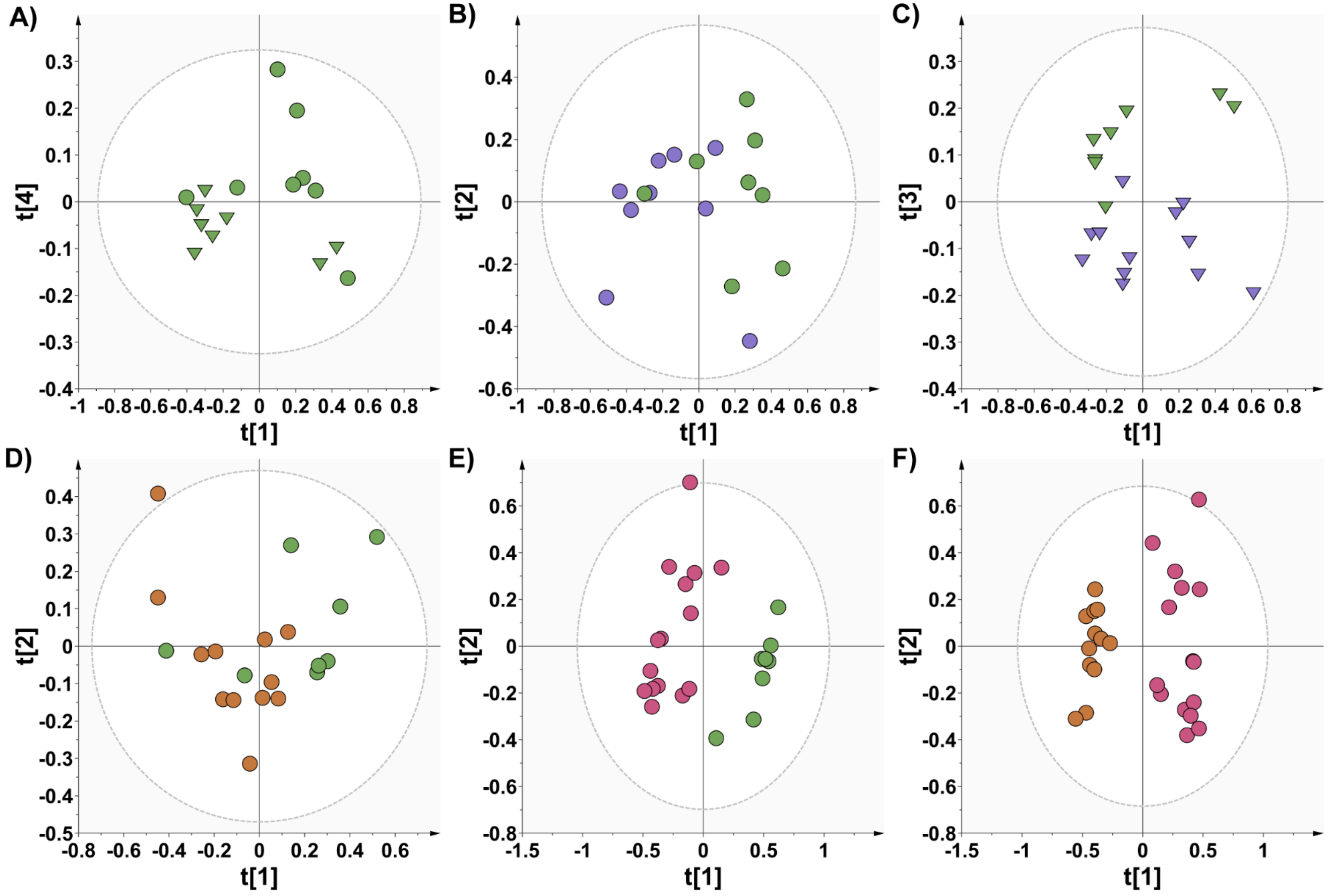


**Supplementary Figure S8: PCA scores plots of pairwise comparisons** A) Uninfected *Drosophila* at 5 dpi (circles) and 9 dpi (inverted triangles). B) *w*Mel-infected *Drosophila* at 5 dpi (purple) and uninfected *Drosophila* at 5 dpi (green). C) *w*Mel-infected *Drosophila* at 9 dpi (purple) and uninfected *Drosophila* at 9 dpi (green). D) Co-infected *Drosophila* at 5 dpi (orange) and uninfected *Drosophila* at 5 dpi (green). E) DCV-infected *Drosophila* at 5 dpi (pink) and uninfected *Drosophila* at 5 dpi (green). F) Co-infected *Drosophila* at 5 dpi (orange) and DCV-infected *Drosophila* at 5 dpi (pink).

**Supplementary Table S1:** Figures of Merit of the Multivariate Statistical Models

| Model [a] | Type [b] | Comparison | A [c] | N [d] | $R^2X$ [e] | $R^2Y$ [e] | $Q^2$ [e] | CV-ANOVA [f] |
|---|---|---|---|---|---|---|---|---|
| M1 | PCA | All cohorts | 13 | 27 | 0.906 | N/A | 0.793 | N/A |
| M2 | PCA | All (excludes QCs) | 13 | 22 | 0.907 | N/A | 0.791 | N/A |
| M3 | PCA | [g][4] -W-V@5 *v.* [7] -W-V@9 | 4 | 21 | 0.779 | N/A | 0.501 | N/A |
| M4 | PCA | [3] +W-V@5 *v.* [6] +W-V@9 | 3 | 19 | 0.654 | N/A | 0.342 | N/A |
| M5 | PCA | [1] +W+V@5 *v.* [5] +W+V@9 | 6 | 44 | 0.815 | N/A | 0.459 | N/A |
| M6 | PCA | [3] +W-V@5 *v.* [4] -W-V@5 | 3 | 71 | 0.694 | N/A | 0.384 | N/A |
| M7 | PCA | [6] +W-V@9 *v.* [7] -W-V@9 | 4 | 16 | 0.731 | N/A | 0.431 | N/A |
| M8 | PCA | [2] -W+V@5 *v.* [4] -W-V@5 | 4 | 21 | 0.827 | N/A | 0.709 | N/A |
| M9 | PCA | [1] +W+V@5 *v.* [2] -W+V@5 | 5 | 19 | 0.853 | N/A | 0.721 | N/A |
| M10 | PCA | [1] +W+V@5 *v.* [3] +W-V@5 | 4 | 20 | 0.7 | N/A | 0.413 | N/A |
| M11 | PCA | [5] +W+V@9 *v.* [6] +W-V@9 | 9 | 23 | 0.913 | N/A | 0.503 | N/A |
| M12 | PCA | [1] +W+V@5 *v.* [4] -W-V@5 | 2 | 20 | 0.506 | N/A | 0.207 | N/A |
| M13 | PCA | [5] +W+V@9 *v.* [7] -W-V@9 | 4 | 15 | 0.775 | N/A | 0.447 | N/A |
| M14 | PCA | Four-way comparison of cohorts at 5 dpi | 5 | 27 | 0.814 | N/A | 0.714 | N/A |
| M15 | PCA | All *w*Mel cohorts | 12 | 71 | 0.89 | N/A | 0.608 | N/A |
| M16 | PLS | [4] -W-V@5 *v.* [7] -W-V@9 | 3 | 16 | 0.672 | 0.888 | 0.345 | 0.533 |
| M17 | PLS | [3] +W-V@5 *v.* [4] -W-V@5 | 5 | 17 | 0.769 | 0.987 | 0.628 | 0.546 |
| M18 | PLS | [6] +W-V@9 *v.* [7] -W-V@9 | 3 | 20 | 0.619 | 0.943 | 0.698 | 0.025 |
| M19 | PLS | [2] -W+V@5 *v.* [4] -W-V@5 | 3 | 23 | 0.746 | 0.965 | 0.875 | $1.653\times10^{-6}$ |
| M20 | PLS | [1] +W+V@5 *v.* [2] -W+V@5 | 2 | 27 | 0.55 | 0.966 | 0.911 | $9.199\times10^{-11}$ |

[a] All model used the same number of x variables (i.e., the number of spectral regions defined as buckets), and all models used pareto scaling
[b] The type of analysis used for the model
[c] This is the number of components
[d] Number of samples
[e] Goodness of fit and predictive ability
[f] p-value of the cross-validated ANOVA (CV-ANOVA)
[g] Numbers between the square brackets denote the meta-data, numbered identity of the cohorts, as seen in the manuscript's methods: Multivariate Statistical Analysis

**Supplementary Table S2**: Concentration Changes in Metabolites for Different Comparisons

| Metabolite | $\frac{-W-V@5}{-W-V@9}$ | $\frac{+W-V@5}{-W-V@5}$ | $\frac{-W+V@5}{-W-V@5}$ | $\frac{+W+V@5}{-W+V@5}$ | $\frac{+W-V@9}{-W-V@9}$ | $\frac{+W+V@5}{+W-V@5}$ | $\frac{+W+V@9}{+W-V@9}$ |
|---|---|---|---|---|---|---|---|
| 2-hydroxyisobutyrate | ⇡ | ⇣ | ↓↓↓ | ↑↑↑ | ⇣ | ⇣ | |
| 2-oxoglutarate | ↑↑↑ | ↓↓↓ | ↓↓↓ | ⇡ | ⇡ | ⇡ | |
| acetate | ⇡ | ⇣ | ⇣ | ⇡ | ↑↑ | | ⇡ |
| acetonitrile | ↑↑↑ | ↓↓ | ⇡ | ⇣ | ⇡ | ⇡ | |
| adenosine | ⇡ | ⇡ | ⇣ | ⇣ | ⇡ | ⇣ | ⇣ |
| alanine | ⇡ | ⇣ | ↓↓↓ | ↑↑↑ | ⇡ | | ⇣ |
| arginine | ⇡ | ⇣ | ⇡ | ⇣ | ⇡ | ⇡ | ⇡ |
| asparagine | ↑↑↑ | ⇣ | ↓↓↓ | ⇡ | ⇡ | ⇣ | ⇡ |
| β-alanine | ↑↑↑ | ⇣ | ↓↓↓ | ↑↑↑ | ⇡ | | |
| dimethylamine | ↑↑↑ | ⇣ | ⇣ | ⇣ | ⇡ | ⇡ | ⇡ |
| formate | ⇡ | | ⇣ | ⇡ | ⇡ | ⇣ | ⇣ |
| fumarate | < LOQ | < LOQ | ↑↑↑ | ↓↓↓ | < LOQ | < LOQ | < LOQ |
| gluconate | ⇡ | ⇡ | ↑↑↑ | ↓↓↓ | ⇡ | ⇡ | ⇡ |
| glucose | ⇡ | ⇣ | ⇡ | ↓↓↓ | ⇡ | ⇡ | |
| glutamate and glutamine | ↑↑↑ | ⇣ | ⇣ | ⇣ | ⇡ | | ⇡ |
| glutamate and proline | ⇡ | ⇡ | ⇣ | ⇡ | ↑↑ | | ⇡ |
| glutamine | ↑↑↑ | ↓↓ | ⇣ | ⇣ | ⇡ | | ⇡ |
| glutamine (excl. oxoglutarate) | ↑↑↑ | ⇣ | ⇣ | ⇣ | ⇡ | | ⇡ |
| glycine | ↑↑↑ | ⇣ | ⇣ | ⇡ | ↑↑ | | ⇣ |
| histidine | ↑↑↑ | ⇣ | ⇣ | | ⇡ | ⇣ | |
| inosine | ↑↑↑ | ⇣ | ↓↓↓ | ⇡ | ⇡ | | ⇡ |
| lactate | ⇡ | ⇣ | ⇣ | ⇡ | ↑↑↑ | ⇡ | ⇡ |
| lipid $(CH_2)_n$ | ⇡ | ⇣ | ⇣ | ⇡ | ⇡ | ⇡ | ⇡ |
| lipid $CH_3$ | ↑↑↑ | ⇣ | ⇣ | ⇡ | ⇡ | ⇡ | ⇡ |
| Lysine | ⇡ | ⇣ | ⇡ | ↓↓↓ | ⇡ | ⇡ | ⇡ |

Continued on next page

| Metabolite | $\frac{-W - V@5}{-W - V@9}$ | $\frac{+W - V@5}{-W - V@5}$ | $\frac{-W + V@5}{-W - V@5}$ | $\frac{+W + V@5}{-W + V@5}$ | $\frac{+W - V@9}{-W - V@9}$ | $\frac{+W + V@5}{+W - V@5}$ | $\frac{+W + V@9}{+W - V@9}$ |
|---|---|---|---|---|---|---|---|
| maltose | ↑↑↑ | ⇣ | ↓↓↓ | ⇡ | ⇡ | | ⇣ |
| NAD+ | ⇡ | ⇣ | ⇣ | ⇡ | ⇡ | | ⇣ |
| o-phosphocholine | ↑↑↑ | ⇣ | | ⇣ | ↑↑ | ⇡ | ⇡ |
| phenylalanine | ⇡ | ⇣ | ⇣ | ⇡ | ↑↑ | | |
| proline | ⇡ | ⇡ | ⇣ | ⇡ | ↑↑ | | ⇣ |
| propionate | ⇡ | ⇣ | ⇡ | ↓↓↓ | ⇡ | ⇡ | ⇡ |
| pyruvate | ⇡ | ⇣ | ↓↓↓ | ↑↑↑ | ⇡ | | ⇣ |
| tyrosine | ⇡ | ⇡ | ↓↓↓ | ↑↑ | ⇡ | | |
| unknown 1 | ⇣ | ⇡ | ↓↓↓ | ↑↑↑ | ⇣ | ⇣ | ⇡ |
| unknown 2 | < LOQ | < LOQ | ↑↑↑ | ↓↓↓ | < LOQ | < LOQ | ⇡ |
| unknown 5 | ⇡ | ⇣ | ↓↓↓ | ↑↑↑ | ⇡ | ⇣ | ⇣ |
| unknown 6 | ⇡ | ⇣ | ↓↓↓ | ↑↑↑ | ⇡ | ⇣ | ⇣ |
| unknown 7 | ⇡ | ⇣ | ↑↑↑ | ↓↓↓ | ⇡ | ⇡ | ⇡ |
| unknown 8 | ↑↑↑ | ⇣ | ⇣ | ⇡ | ↑↑ | | ⇣ |
| uracil | ↑↑↑ | ⇣ | ⇣ | | ⇡ | ⇣ | ⇡ |
| uridine | ⇡ | ⇣ | ⇣ | ⇡ | ⇡ | ⇣ | ⇣ |
| valine | ⇡ | ⇣ | ↑↑↑ | ↓↓↓ | ↑↑ | ⇡ | ⇡ |

Note: Arrows facing upwards indicate metabolites that were determined to be at higher levels in one cohort relative to the cohort listed below (e.g., cohort under study/relative cohort). Arrows facing downwards indicate metabolites that were determined to be at lower levels in one cohort relative to another. ⇡ or ⇣: indicates metabolites that were either increased or decreased, with no statistical significance ($p > 0.05$). ↓↓ or ↑↑: indicates metabolites that were either significantly increased or decreased ($p \leq 0.01$). ↓↓↓ or ↑↑↑: indicates metabolites that were either significantly increased or decreased ($p \leq 0.001$). In the comparisons where both cohorts shared the integration average less than limit of quantification were indicated with < LOQ, or the metabolite was excluded if the integration average was less than the limit of quantification for all cohorts.

**Supplementary Table S3:** List of Primers used for RT-PCR

| Primer Name [a] | Sequence (5'→3') | Amplicon Fragment Size (bp) |
|---|---|---|
| DCV-rt-fw | AGG CTG TGT TTG CGC GAA G | 203 |
| DCV-rt-rv | AAT GGC AAG CGC ACA CAA TTA | |
| FHV-rt-fw | AGT GGT CAG CCG AAA GGA TG | 546 |
| FHV-rt-rv | CCC GTA GAA CCA AGG GAC AC | |
| CrPV-rt-fw | CAC TCT TCA GGC TGA TGG CA | 199 |
| CrPV-rt-rv | CTT CCG GGG AGT TCC TGT TC | |
| SINV-rt-fw | GCG TAC GTC GAA TTG TCA GC | 200 |
| SINV-rt-rv | GTC TTC TTC CCA CAC AGC GA | |
| RpL32-fw | GAC GCT TCA AGG GAC AGT ATC TG | 143 |
| RpL32-rv | AAA CGC GGT TCT GCA TGA G | |

[a] Forward primer (fw); reverse primer (rv); [Flock] House virus (FHV); Cricket paralysis virus (CrPV); Sindbis virus (SINV)

**Supplementary Table 4:** Primers to confirm absence/presence of *w*Mel infection.

| Primer | Sequence (5’→ 3’) |
|---|---|
| RpL32-fw [a] | GAC GCT TCA AGG GAC AGT ATC TG |
| RpL32-rv [b] | AAA CGC GGT TCT GCA TGA G |
| *wsp*-fw | TGG TCC AAT AAG TGA TGA AGA AAC |
| *wsp*-rv | AAA AAT TAA ACG CTA CTC CA |

[a] Forward primer (fw)
[b] Reverse primer (rv)

**Supplementary Table S5:** PCR experiments to confirm absence/presence of *w*Mel.

| Template | Primer | Purpose |
|---|---|---|
| *w*Mel-free samples | *wsp* | To confirm the presence or absence of *w*Mel |
| *w*Mel samples | *wsp* | |
| *w*Mel samples | RpL32 | Endogenous positive control confirming the presence of *D. melanogaster* genomic DNA |
| *w*Mel-free samples | RpL32 | |
| No template control | *wsp* | Demonstrate absence of genomic contamination |
| No template control | RpL32 | |
| Negative control [a] | *wsp* | To confirm the validity of the confirmation of *w*Mel absence or presence |
| Negative control | RpL32 | |
| Positive control [b] | *wsp* | |
| Positive control | RpL32 | |

[c] Control consists of DNA extracted from *Drosophila* previously validated to be *w*Mel-free

[d] Control consists of DNA extracted from *Drosophila* previously validated to be *w*Mel-infected

**Supplementary Table S6:** Key metabolites and associated observed NMR resonances.

| Metabolite | $^{1}H$ Peak(s)/$^{13}C$ Peak(s) |
|---|---|
| 2-Hydroxyisobutyrate | 1.3585(s)/28.149 |
| 2-Oxoglutarate | 2.9965(t)/38.515; 2.4305(t)/ [b] |
| Acetate | 1.9105(s)/ 26.002 |
| Acetonitrile | 2.0645(s)/ 3.57 |
| Adenosine | 8.4315(s)/ [b]; 8.2555(s)/ [b]; 6.0635(d)/ [b] |
| Alanine | 1.4755(d)/18.951 |
| Arginine | 1.908(m)/30.318; 1.7175(m)/26.553; 1.6545(m)/ 26.516 |
| Asparagine | 2.9285(dd)/ [b]; 2.8705 (dd)/ [b] |
| β-Alanine | 3.1725(t)/39.392; 2.5565(t)/36.388 |
| Dimethylamine | 2.7205(s)/ 41.422 |
| Formate | 8.4465(s) [b] |
| Fumarate | 6.5105(s) [b] |
| Gluconate | 4.1295(d)/76.838; 4.0285(t)/ 73.753 |
| Glucose | 5.2315(d)/94.779; 4.6455(d)/98.622; 3.8985 (dd)/63.446; 3.8275(m)/ [b]; 3.7245(q)/ [b]; 3.7055(t); 75.562; 3.5325(dd)/74.273; 3.4825(t)/78.618; 3.4475(m)/78.618; 3.4085(t)/72.264; 3.3965(t)/ [b]; 3.2365(dd)/76.967 |
| Glutamate and Glutamine | 2.1285(m)/29.743 |
| Glutamate and Proline | 2.3265(m)/G: 36.164[c] P: 31.682[c]; 2.0805(m)/G:29.752 P:31.638; |
| Glutamine | 2.442305(m)/33.356 |
| Glycine | 3.5545(s)/44.268 |
| Guanosine | 7.9935(s)/ [b]; 5.8945(d)/ [b] |
| histidine | 7.9175(s)/138.559;7.098(s)/119.787 |
| inosine | 8.339(s)/ [b]; 8.23(s)/ [b]; 6.0925(d)/ [b] |
| lactate | 4.1115(q)/71.261; 1.3235(d)/22.767 |
| lipid $(CH_2)_n$ | 1.2775/32.047 |
| lipid $CH_3$ | 0.8715/16.495 |
| lysine | 3.0185(t)/42.046 |
| maltose | 5.4035(2×d)/102.209; 4.6575(d)/98.622; 3.9655(t)/75.562; 3.8985(dd)/63.446; 3.6345(2×t)/ [b] |

Continued on the next page

| Metabolites | [1]H Peak/s |
|---|---|
| NAD | 9.3295(s)/ [b]; 9.1365/ [b], 9.1285(d)/ [b]; 8.8305(d)/ [b]; 8.4185(s)/ [b]; 8.1675(s)/ [b]; 6.0315(d)/ [b] |
| NADP | 9.2875(s)/ [b]; 9.0915(d)/ [b]; 8.8045(d)/ [b]; 8.4065(s)/ [b];6.0785(d)/ [b] |
| o-phosphocholine | 3.2095(s)/56.653 |
| phenylalanine | 7.4195(t)/ [b]; 7.3675(t)/ [b]; 7.3245(d)/ [b] |
| proline | 3.403(m)/48.881; 1.994(m)/26.443 |
| propionate | 2.1685(q)/33.465; 1.0456(t)/12.941 |
| pyruvate | 2.3645(s)/29.171 |
| tyrosine | 7.1885(d)/ [b]; 6.8975(d)/ [b] |
| unknown 1 | 1.5375(d)/20.668; 1.5245(d)/19.437 |
| unknown 2 | 1.1215(m)/ [b] |
| unknown 3 | 2.6445(d)/48.282 |
| unknown 4 | 2.7445(s)/39.215 |
| unknown 5 | 1.6305(s)/ [b] |
| unknown 6 | 1.7895(s)/ [b] |
| unknown 7 | 4.2995(d)/ [b] |
| unknown 8 | 4.4635(d)/ [b] |
| uracil | 7.5375(d)/ [b]; 5.7995(d)/ [b] |
| uridine | 7.8625(d)/ [b]; 5.9095(d)/ [b]; 5.8995(d)/ [b] |
| valine | 1.0365(d)/20.668; 0.9835(d)/19.437 |

[a] s, singlet; d, doublet; dd, doublet of a doublet; t, triplet; m, multiplet

[b] Could not unambiguously confirm carbon peak

[c] This indicates while there is overlap in the proton spectra for these metabolites, in the 2D spectra, the carbon peaks distinguish out the metabolites from one another where (G) is denoting the carbon peaks of glutamine and (P) denotes the carbon peaks of proline for these corresponding proton signals

**Supplementary Table S7:** NMR Spectra Integration Regions

| Metabolite | Integration region/s (ppm) |
|---|---|
| 2-hydroxyisobutyrate | (1.36346 - 1.35475) |
| 2-oxoglutarate | (3.001 - 2.98404) |
| acetate | (1.91623 - 1.90447) |
| acetonitrile | (2.06734 - 2.06214) |
| adenosine | (8.25723 - 8.24943) + (6.07532 - 6.05683) |
| alanine | (1.4834 - 1.45635) |
| arginine | (1.75489 - 1.68415) |
| asparagine | (2.95883 - 2.94539) + (2.94034 - 2.92002) |
| β-alanine | (3.18709 - 3.16127) + (2.56175 - 2.53318) |
| dimethylamine | (2.72797 - 2.71163) |
| formate | (8.45264 - 8.4398) |
| fumarate | (6.5242 - 6.4967) |
| gluconate | (4.03672 - 4.01854) |
| glucose | (5.2399 - 5.21606) + (4.65275 - 4.62525) + (3.90716 - 3.87844) + (3.54506 - 3.51298) + (3.50198 - 3.47738) |
| glutamate and glutamine | (2.15183 - 2.0872) |
| glutamate and proline | (2.3584 - 2.31103) + (2.05756 - 2.01661) |
| glutamine and oxoglutarate | (2.48398 - 2.40071) |
| glutamine (excl. oxoglutarate) | (2.48398 - 2.46000) |
| glycine | (3.55576 - 3.54751) |
| guanosine | (5.893 - 5.88098) |
| histidine | (7.924 - 7.90888) + (7.10508 - 7.08033) |
| inosine | (8.234 - 8.22774) + (6.09854 - 6.08189) + (4.4442 - 4.41624) |
| lactate | (1.32863 - 1.31121) |
| lipid $(CH_2)_n$ | (1.30815 - 1.21954) |
| lipid $CH_3$ | (0.903734 - 0.767757) |
| lysine | (3.03156 - 3.01124) |
| maltose | (5.41575 - 5.39971) + (4.66451 - 4.65229) |
| NAD | (9.33588 - 9.32381) + (9.14337 - 9.12076) + (8.83628 - 8.81153) + (8.42376 - 8.41169) + (6.03452 - 6.01848) |
| NADP | (9.30364 - 9.27232) + (9.10334 - 9.0815) + (8.14432 - 8.13454) |
| o-phosphocholine | (3.21306 - 3.20313) |
| phenylalanine | (7.43754 - 7.39889) + (7.38727 - 7.35198) + (7.33502 - 7.30248) |
| proline | (3.34186 - 3.30916) |
| propionate | (2.19277 - 2.15137) + (1.06095 - 1.04048) |
| pyruvate | (2.37046 - 2.35778) |
| tyrosine | (7.194 - 7.17124) + (6.90203 - 6.87973) |
| unknown 1 | (1.54161 - 1.50968) |
| unknown 2 | (1.14269 - 1.09914) |
| unknown 3 | (2.67236 - 2.63615) |
| unknown 4 | (2.75273 - 2.73577) |
| unknown 5 | (1.63465 - 1.62549) |
| unknown 6 | (1.79615 - 1.77919) |
| unknown 7 | (4.3073 - 4.29584) |
| unknown 8 | (4.45856 - 4.44741) |

Continued on the next page

| Metabolites | Integration region/s (ppm) |
| --- | --- |
| uracil | (7.5451 - 7.52356) + (5.80657 - 5.78243) |
| uridine | (7.87618 - 7.85235) + (5.91459 - 5.88021) + (5.89198 - 5.8825) |
| valine | (1.0307 - 1.02107) + (0.989904 - 0.968056) |

**Supplementary Table S8:** Metabolite Average Integration for the Cohorts Collected at 5 dpi.

| Metabolite | +W+V@5 | -W+V@5 | +W-V@5 | -W-V@5 |
|---|---|---|---|---|
| 2-hydroxyisobutyrate | $(4.632 \pm 2.706) \times 10^6$ | < LOQ | $(7.138 \pm 2.220) \times 10^6$ | $(8.721 \pm 3.307) \times 10^6$ |
| 2-oxoglutarate | $(2.43 \pm 0.992) \times 10^7$ | $(2.118 \pm 0.563) \times 10^7$ | $(2.240 \pm 0.620) \times 10^7$ | $(3.548 \pm 0.439) \times 10^7$ |
| acetate | $(7.039 \pm 2.776) \times 10^7$ | $(6.888 \pm 3.535) \times 10^7$ | $(7.016 \pm 2.055) \times 10^7$ | $(8.716 \pm 3.718) \times 10^7$ |
| acetonitrile | $(6.524 \pm 4.705) \times 10^7$ | $(1.103 \pm 1.092) \times 10^8$ | $(2.296 \pm 2.205) \times 10^7$ | $(7.413 \pm 1.919) \times 10^7$ |
| adenosine | $(1.019 \pm 0.461) \times 10^7$ | $(1.204 \pm 0.685) \times 10^7$ | $(1.545 \pm 0.750) \times 10^7$ | $(1.504 \pm 0.670) \times 10^7$ |
| alanine | $(5.01 \pm 1.619) \times 10^8$ | $(2.922 \pm 0.799) \times 10^8$ | $(4.931 \pm 1.289) \times 10^8$ | $(5.773 \pm 0.656) \times 10^8$ |
| arginine | $(7.123 \pm 2.500) \times 10^7$ | $(9.644 \pm 2.716) \times 10^7$ | $(6.495 \pm 1.560) \times 10^7$ | $(7.967 \pm 1.487) \times 10^7$ |
| asparagine | $(9.173 \pm 2.793) \times 10^6$ | $(5.396 \pm 1.580) \times 10^6$ | $(8.969 \pm 1.687) \times 10^6$ | $(1.167 \pm 0.185) \times 10^7$ |
| β-alanine | $(5.325 \pm 1.959) \times 10^8$ | $(2.876 \pm 0.761) \times 10^8$ | $(5.293 \pm 1.489) \times 10^8$ | $(6.048 \pm 0.373) \times 10^8$ |
| dimethylamine | $(2.503 \pm 1.733) \times 10^7$ | $(2.786 \pm 2.165) \times 10^7$ | $(1.638 \pm 0.958) \times 10^7$ | $(3.711 \pm 0.548) \times 10^7$ |
| formate | $(2.002 \pm 0.415) \times 10^6$ | $(1.680 \pm 0.350) \times 10^6$ | $(2.233 \pm 0.322) \times 10^6$ | $(2.218 \pm 0.364) \times 10^6$ |
| fumarate | < LOQ | $(8.802 \pm 2.639) \times 10^6$ | < LOQ | < LOQ |
| gluconate | $(6.847 \pm 5.110) \times 10^7$ | $(3.306 \pm 1.359) \times 10^8$ | $(4.275 \pm 1.856) \times 10^7$ | $(4.615 \pm 4.707) \times 10^7$ |
| glucose | $(1.712 \pm 0.580) \times 10^9$ | $(3.628 \pm 1.283) \times 10^9$ | $(1.611 \pm 0.414) \times 10^9$ | $(2.123 \pm 0.252) \times 10^9$ |
| glutamate and glutamine | $(1.188 \pm 0.486) \times 10^8$ | $(1.439 \pm 0.437) \times 10^8$ | $(1.187 \pm 0.322) \times 10^8$ | $(1.603 \pm 0.192) \times 10^8$ |
| glutamate and proline | $(1.971 \pm 0.683) \times 10^8$ | $(1.821 \pm 0.490) \times 10^8$ | $(1.979 \pm 0.583) \times 10^8$ | $(1.921 \pm 0.221) \times 10^8$ |
| glutamine | $(1.033 \pm 0.440) \times 10^8$ | $(1.329 \pm 0.451) \times 10^8$ | $(1.008 \pm 0.290) \times 10^8$ | $(1.558 \pm 0.219) \times 10^8$ |
| glutamine (excl. oxoglutarate) | $(1.173 \pm 0.559) \times 10^7$ | $(1.492 \pm 0.507) \times 10^7$ | $(1.157 \pm 0.400) \times 10^7$ | $(1.734 \pm 0.393) \times 10^7$ |
| glycine | $(6.360 \pm 2.060) \times 10^7$ | $(5.256 \pm 1.387) \times 10^7$ | $(6.487 \pm 1.739) \times 10^7$ | $(6.84 \pm 1.127) \times 10^7$ |
| guanosine | < LOQ | < LOQ | < LOQ | < LOQ |
| histidine | $(8.865 \pm 3.181) \times 10^7$ | $(8.941 \pm 2.375) \times 10^7$ | $(9.435 \pm 2.3950 \times 10^7$ | $(1.179 \pm 0.128) \times 10^8$ |
| inosine | $(6.930 \pm 1.770) \times 10^7$ | $(5.088 \pm 1.495) \times 10^7$ | $(6.649 \pm 1.629) \times 10^7$ | $(8.266 \pm 2.152) \times 10^7$ |
| lactate | $(1.177 \pm 0.371) \times 10^8$ | $(9.301 \pm 3.198) \times 10^7$ | $(1.128 \pm 0.363) \times 10^8$ | $(1.161 \pm 0.167) \times 10^8$ |
| lipid $(CH_2)_n$ | $(2.005 \pm 0.715) \times 10^8$ | $(1.361 \pm 0.464) \times 10^8$ | $(1.656 \pm 0.697) \times 10^8$ | $(2.145 \pm 0.506) \times 10^8$ |
| lipid $CH_3$ | $(1.900 \pm 0.592) \times 10^8$ | $(1.582 \pm 0.368) \times 10^8$ | $(1.735 \pm 0.436) \times 10^8$ | $(2.075 \pm 0.238) \times 10^8$ |
| lysine | $(1.710 \pm 0.668) \times 10^7$ | $(3.471 \pm 1.204) \times 10^7$ | $(1.509 \pm 0.459) \times 10^7$ | $(2.096 \pm 0.559) \times 10^7$ |
| maltose | $(2.842 \pm 1.082) \times 10^8$ | $(2.017 \pm 0.598) \times 10^8$ | $(2.910 \pm 0.733) \times 10^8$ | $(3.270 \pm 0.283) \times 10^8$ |
| NAD | $(1.488 \pm 0.730) \times 10^7$ | $(1.181 \pm 0.314) \times 10^7$ | $(1.509 \pm 0.546) \times 10^7$ | $(1.792 \pm 0.333) \times 10^7$ |

Continued on the next page

| Metabolite | +W+V@5 | -W+V@5 | +W-V@5 | -W-V@5 |
|---|---|---|---|---|
| NADP | < LOQ | < LOQ | < LOQ | < LOQ |
| o-phosphocholine | $(4.363 \pm 1.525) \times 10^{8}$ | $(4.700 \pm 1.180) \times 10^{8}$ | $(4.128 \pm 1.050) \times 10^{8}$ | $(4.713 \pm 0.853) \times 10^{8}$ |
| phenylalanine | $(1.355 \pm 0.394) \times 10^{7}$ | < LOQ | $(1.350 \pm 0.293) \times 10^{7}$ | $(1.455 \pm 0.264) \times 10^{7}$ |
| proline | $(7.039 \pm 2.314) \times 10^{7}$ | $(5.002 \pm 1.311) \times 10^{7}$ | $(7.339 \pm 2.141) \times 10^{7}$ | $(6.441 \pm 0.469) \times 10^{7}$ |
| propionate | $(2.446 \pm 1.175) \times 10^{7}$ | $(8.740 \pm 5.064) \times 10^{7}$ | $(2.103 \pm 0.618) \times 10^{7}$ | $(3.385 \pm 2.480) \times 10^{7}$ |
| pyruvate | $(2.297 \pm 1.144) \times 10^{8}$ | $(5.871 \pm 3.588) \times 10^{7}$ | $(2.130 \pm 0.607) \times 10^{8}$ | $(3.350 \pm 1.043) \times 10^{8}$ |
| tyrosine | $(7.577 \pm 2.719) \times 10^{6}$ | < LOQ | $(7.985 \pm 2.249) \times 10^{6}$ | $(7.449 \pm 1.171) \times 10^{6}$ |
| unknown 1 | $(9.359 \pm 5.027) \times 10^{6}$ | < LOQ | $(1.937 \pm 0.737) \times 10^{7}$ | $(1.593 \pm 0.609) \times 10^{7}$ |
| unknown 2 | < LOQ | $(3.212 \pm 1.503) \times 10^{7}$ | < LOQ | < LOQ |
| unknown 3 | $(2.197 \pm 1.038) \times 10^{7}$ | $(4.566 \pm 1.344) \times 10^{7}$ | $(1.817 \pm 0.690) \times 10^{7}$ | $(2.263 \pm 0.660) \times 10^{7}$ |
| unknown 4 | $(3.589 \pm 1.416) \times 10^{7}$ | $(5.325 \pm 1.583) \times 10^{7}$ | $(4.054 \pm 0.793) \times 10^{7}$ | $(6.292 \pm 1.310) \times 10^{7}$ |
| unknown 5 | $(1.676 \pm 0.411) \times 10^{7}$ | $(6.445 \pm 1.723) \times 10^{6}$ | $(1.903 \pm 0.404) \times 10^{7}$ | $(1.990 \pm 0.280) \times 10^{7}$ |
| unknown 6 | $(5.478 \pm 1.451) \times 10^{6}$ | < LOQ | $(6.206 \pm 1.415) \times 10^{6}$ | $(6.806 \pm 1.307) \times 10^{6}$ |
| unknown 7 | $(2.376 \pm 0.834) \times 10^{6}$ | $(8.919 \pm 2.729) \times 10^{6}$ | $(1.441 \pm 0.648) \times 10^{6}$ | $(2.410 \pm 0.978) \times 10^{6}$ |
| unknown 8 | $(3.020 \pm 1.068) \times 10^{7}$ | $(2.162 \pm 0.582) \times 10^{7}$ | $(3.014 \pm 0.793) \times 10^{7}$ | $(3.087 \pm 0.428) \times 10^{7}$ |
| uracil | $(5.287 \pm 1.363) \times 10^{6}$ | $(5.251 \pm 1.381) \times 10^{6}$ | $(5.662 \pm 1.704) \times 10^{6}$ | $(6.346 \pm 0.903) \times 10^{6}$ |
| uridine | $(8.214 \pm 2.588) \times 10^{6}$ | < LOQ | $(9.046 \pm 2.227) \times 10^{6}$ | $(9.742 \pm 2.097) \times 10^{6}$ |
| valine | $(1.934 \pm 0.614) \times 10^{7}$ | $(3.947 \pm 1.192) \times 10^{7}$ | $(1.783 \pm 0.471) \times 10^{7}$ | $(2.015 \pm 0.930) \times 10^{7}$ |

Note: The metabolite integration is given ± the standard deviation. Any of the values that were below the limit of quantification (< LOQ) are identified as such. The designations read as *w*Mel-infected (+W), *w*Mel-free (-W), DCV-infected (+V), DCV-free (-V), at 5 dpi (@5), and at 9 dpi (@9).

**Supplementary Table S9:** Metabolite Average Integration for the Cohorts Collected at 9 dpi.

| Metabolite | +W+V@9 | +W-V@9 | -W-V@9 |
|---|---|---|---|
| 2-hydroxyisobutyrate | (4.027 ± 3.272) $\times 10^6$ | (4.065 ± 1.431) $\times 10^6$ | (6.077 ± 2.529) $\times 10^6$ |
| 2-oxoglutarate | (2.438 ± 0.757) $\times 10^7$ | (2.418 ± 0.856) $\times 10^7$ | (1.853 ± 0.502) $\times 10^7$ |
| acetate | (1.124 ± 0.522) $\times 10^8$ | (9.214 ± 3.032) $\times 10^7$ | (5.073 ± 1.032) $\times 10^7$ |
| acetonitrile | (6.131 ± 4.159) $\times 10^7$ | (5.888 ± 4.677) $\times 10^7$ | (2.025 ± 1.676) $\times 10^7$ |
| adenosine | (9.188 ± 8.161) $\times 10^6$ | (1.964 ± 0.940) $\times 10^7$ | (1.395 ± 0.514) $\times 10^7$ |
| alanine | (5.287 ± 1.691) $\times 10^8$ | (5.758 ± 1.716) $\times 10^8$ | (3.595 ± 0.734) $\times 10^8$ |
| arginine | (9.352 ± 3.285) $\times 10^7$ | (7.383 ± 2.659) $\times 10^7$ | (4.414 ± 0.763) $\times 10^7$ |
| asparagine | (9.689 ± 2.580) $\times 10^6$ | (8.818 ± 3.124) $\times 10^6$ | (6.922 ± 1.437) $\times 10^6$ |
| β-alanine | (5.190 ± 1.644) $\times 10^8$ | (5.398 ± 1.667) $\times 10^8$ | (3.705 ± 0.689) $\times 10^8$ |
| dimethylamine | (3.214 ± 1.302) $\times 10^7$ | (2.566 ± 1.749) $\times 10^7$ | (8.892 ± 1.366) $\times 10^6$ |
| formate | (2.307 ± 0.544) $\times 10^6$ | (2.401 ± 0.653) $\times 10^6$ | (2.015 ± 0.241) $\times 10^6$ |
| fumarate | < LOQ | < LOQ | < LOQ |
| gluconate | (1.167 ± 0.785) $\times 10^8$ | (6.401 ± 5.488) $\times 10^7$ | (2.608 ± 0.847) $\times 10^7$ |
| glucose | (1.984 ± 0.742) $\times 10^9$ | (2.002 ± 0.824) $\times 10^9$ | (1.260 ± 0.270) $\times 10^9$ |
| glutamate and glutamine | (1.300 ± 0.396) $\times 10^8$ | (1.158 ± 0.415) $\times 10^8$ | (9.322 ± 2.102) $\times 10^7$ |
| glutamate and proline | (2.183 ± 0.701) $\times 10^8$ | (2.061 ± 0.648) $\times 10^8$ | (1.211 ± 0.232) $\times 10^8$ |
| glutamine | (1.049 ± 0.316) $\times 10^8$ | (9.898 ± 3.436) $\times 10^7$ | (8.681 ± 2.422) $\times 10^7$ |
| glutamine (excl. oxoglutarate) | (1.183 ± 0.431) $\times 10^7$ | (1.078 ± 0.431) $\times 10^7$ | (1.004 ± 0.269) $\times 10^7$ |
| glycine | (7.224 ± 2.387) $\times 10^7$ | (7.488 ± 2.370) ×107 | (4.016 ± 0.845) ×107 |
| guanosine | < LOQ | < LOQ | < LOQ |
| histidine | (1.038 ± 0.356) $\times 10^8$ | (1.023 ± 0.323) $\times 10^8$ | (6.932 ± 1.415) $\times 10^7$ |
| inosine | (8.641 ± 3.360) $\times 10^7$ | (6.863 ± 2.097) $\times 10^7$ | (4.669 ± 0.884) $\times 10^7$ |
| lactate | (1.537 ± 0.696) $\times 10^8$ | (1.457 ± 0.439) $\times 10^8$ | (7.542 ± 1.883) $\times 10^7$ |
| lipid $(CH_2)_n$ | (2.272 ± 1.494) $\times 10^8$ | (1.879 ± 0.880) $\times 10^8$ | (1.469 ± 0.647) $\times 10^8$ |
| lipid $CH_3$ | (2.237 ± 0.713) $\times 10^8$ | (1.906 ± 0.659) $\times 10^8$ | (1.280 ± 0.270) $\times 10^8$ |
| lysine | (2.515 ± 0.710) $\times 10^7$ | (1.891 ± 0.889) $\times 10^7$ | (1.321 ± 0.155) $\times 10^7$ |
| maltose | (2.433 ± 0.678) $\times 10^8$ | (2.950 ± 0.780) $\times 10^8$ | (1.964 ± 0.442) $\times 10^8$ |
| NAD | (1.303 ± 0.591) $\times 10^7$ | (1.409 ± 0.415) $\times 10^7$ | (1.083 ± 0.352) $\times 10^7$ |
| NADP | < LOQ | < LOQ | < LOQ |

Continued on the next page

| Metabolite | +W+V@9 | +W-V@9 | -W-V@9 |
|---|---|---|---|
| o-phosphocholine | $(5.357 \pm 1.853) \times 10^8$ | $(4.512 \pm 1.440) \times 10^8$ | $(2.646 \pm 0.507) \times 10^8$ |
| phenylalanine | $(1.617 \pm 0.606) \times 10^7$ | $(1.580 \pm 0.487) \times 10^7$ | < LOQ |
| proline | $(7.073 \pm 2.279) \times 10^7$ | $(7.690 \pm 2.303) \times 10^7$ | $(4.452 \pm 0.966) \times 10^7$ |
| propionate | $(5.003 \pm 2.677) \times 10^7$ | $(2.841 \pm 2.248) \times 10^7$ | $(1.914 \pm 0.466) \times 10^7$ |
| pyruvate | $(1.535 \pm 0.525) \times 10^8$ | $(2.027 \pm 0.605) \times 10^8$ | $(1.522 \pm 0.387) \times 10^8$ |
| tyrosine | $(8.073 \pm 3.745) \times 10^6$ | $(8.054 \pm 2.251) \times 10^6$ | $(6.463 \pm 1.709) \times 10^6$ |
| unknown 1 | $(1.169 \pm 0.838) \times 10^7$ | $(9.006 \pm 4.079) \times 10^6$ | $(1.857 \pm 0.836) \times 10^7$ |
| unknown 2 | $(5.369 \pm 3.871) \times 10^6$ | < LOQ | < LOQ |
| unknown 3 | $(3.181 \pm 1.146) \times 10^7$ | $(1.844 \pm 0.619) \times 10^7$ | $(1.365 \pm 0.404) \times 10^7$ |
| unknown 4 | $(3.709 \pm 1.683) \times 10^7$ | $(3.734 \pm 1.472) \times 10^7$ | $(3.797 \pm 1.035) \times 10^7$ |
| unknown 5 | $(1.621 \pm 0.460) \times 10^7$ | $(1.843 \pm 0.404) \times 10^7$ | $(1.398 \pm 0.213) \times 10^7$ |
| unknown 6 | $(4.983 \pm 1.778) \times 10^6$ | $(5.996 \pm 1.519) \times 10^6$ | $(4.358 \pm 0.837) \times 10^6$ |
| unknown 7 | $(2.988 \pm 0.714) \times 10^6$ | $(2.288 \pm 2.068) \times 10^6$ | $(1.584 \pm 0.387) \times 10^6$ |
| unknown 8 | $(3.237 \pm 1.093) \times 10^7$ | $(3.306 \pm 1.037) \times 10^7$ | $(1.875 \pm 0.365) \times 10^7$ |
| uracil | $(6.548 \pm 1.516) \times 10^6$ | $(6.281 \pm 1.926) \times 10^6$ | < LOQ |
| uridine | $(8.511 \pm 1.770) \times 10^6$ | $(9.195 \pm 2.713) \times 10^6$ | < LOQ |
| valine | $(3.085 \pm 1.302) \times 10^7$ | $(2.253 \pm 0.953) \times 10^7$ | $(1.117 \pm 0.240) \times 10^7$ |

Note: The metabolite integration is given ± the standard deviation. Any of the values that were below the limit of quantification (< LOQ) are identified as such. The designations read as *w*Mel-infected (+W), *w*Mel-free (-W), DCV-infected (+V), DCV-free (-V), at 5 dpi (@5), and at 9 dpi (@9)

**Dataset S1 (separate file)**

The NMR data presented in this study can be found in the online repository – MetaboLights. It can be found under the accession number ####.
